# Designing a Multi-Epitope Vaccine Candidate against *Salmonella* Typhi Using Immunoinformatics Approach

**DOI:** 10.1101/2022.01.31.478495

**Authors:** Ahmad Salman Sirajee, Sunjukta Ahsan

## Abstract

*Salmonella enterica* subsp. enterica serovar Typhi (*Salmonella* Typhi) is the causative agent of typhoid. A large population living in developing and under-developed countries are mainly susceptible to the infection of this pathogen. The use of an effective vaccine targeting the highest-risk populations could be an extremely useful approach in the prevention of typhoid. The currently licenced and widely used typhoid conjugate vaccine (Vi-TCV) can promote emergence of Vi antigen negative strains, necessitating the design of alternative vaccine candidates. Vaccine consisting of multiple epitopes from different surface proteins can serve as a solution to this problem. In this study, 10 *Salmonella* Typhi whole genome sequences were retrieved from NCBI database. From the proteomes of these organisms, epitopes from four surface proteins such as peptidoglycan-associated lipoprotein (Accession No. AYU85048.1), outer membrane protein S1 (Accession No. AYU86170.1), outer membrane protein F (Accession No. AYU85220.1 and outer membrane porin L (Accession No. AYU87629.1) returned better results than some other surface proteins in terms of antigenicity, promiscuity to interact with MHC alleles, non-allergenicity, non-similarity with human proteins, conservation and population coverage. These epitopes were joined together using suitable adjuvant and linkers, resulting in a vaccine candidate comprising 201 amino acids. This multi epitope vaccine candidate exhibited promising results in molecular docking, molecular dynamics simulation and immune simulation. BLAST search revealed that the chosen epitopes for vaccine development against *Salmonella* Typhi are 100% identical with corresponding epitopes of *Salmonella* Paratyphi A, *Salmonella* Typhimurium and *Salmonella* Enteritidis, indicating that the proposed vaccine might give cross protection against major *Salmonella* pathogenic serovars. The prospective results in this dry laboratory work demonstrates potential positive outcomes in the future for in vivo animal model testing and clinical analyses.

## Introduction

*Salmonella enterica* serovar Typhi (*S*. Typhi) is one of the most notorious foodborne diseases worldwide, most prominently in urban slums and rural areas of low and middle-income countries where access to safe food, pure water and proper sanitation is limited. Particularly, in Asia (Ochiai et. al., 2008) and sub-Saharan Africa (Marks et. al., 2017), this disease is not monitored adequately. Moreover, the confirmation of the diagnosis of Typhoid is difficult (Mogasale et. al., 2016). The severity of this disease depends on the immunity of an individual. Children aging below 5 years, elderly people and immunosuppressed people are more susceptible to Typhoid.

According to 16S rRNA nomenclature system, the genus *Salmonella* is classified into two species, such as *Salmonella enterica* (type species) and *Salmonella bongori*. Among all the subspecies of *Salmonella, S. enterica* subsp. enterica is responsible for about 99% of *Salmonella* infections in humans. In addition, the Kauffman and White scheme classifies *Salmonella* into three serotypes based on their three major antigenic determinants, such as: somatic (O), capsular (K) and flagellar (H) (Brenner et. al., 2000). According to this scheme, the *S*. Typhi is placed in Group D due to having O-antigen type O9-12, phase1 flagellin type H:d, and Vi capsule.

*Salmonella* Typhi can enter into human body upon ingestion of this bacterium through contaminated food or water. The incubation period for this pathogen usually ranges from 8 to 14 days (Wangdi et. al., 2012). This pathogen can enter into long-term carrier status by reaching to the gallbladder, which is an immune privileged site (Gilman et. al., 1979), and also, by surviving within macrophages, granting their access in the reticuloendothelial system (RES) (Monack et. al., 2004).

The relentless emergence of antimicrobial resistant strains of *Salmonella* is making things difficult in combating this pathogen. For example, the recent emergence of extensively drug resistant (XDR) *S*. Typhi strains has made the treatment of typhoid fever tough and expensive (Klemm et. al., 2018). This disease is the deadliest particularly among the children below 2 years of age, which necessitates the early programmatic intervention with vaccination crucial (Jeon et. al., 2018).

The obvious way to lower the occurrence of a foodborne disease to almost zero is the development of infrastructure to maximize the availability of safe water and hygienic environment to everyone. While this a long term and costly process, spreading knowledge of good hygiene practice and making people aware of food safety, in addition to the use of an effective vaccine can be life saver. In case of *S*. Typhi, there are three types of vaccines available that are currently licensed, such as: the live attenuated Ty21a vaccine, the Vi-polysaccharide vaccine and the typhoid conjugate vaccine (TCV) vaccines. However, these vaccines have some limitations in terms of safety and efficacy. For example, the oral vaccine Ty21a uses attenuated *S*. Typhi. If somehow this strain mutates to regain pathogenic characteristics inside human body, it can pose life threatening cases. Also, this vaccine is not safe for an immunocompromised individual. This type of vaccine is not suitable for children aging below 5 years. Vi polysaccharide vaccine is taken parenterally, which may cause uncomfortable pain to the vaccine taker (MacLennan et. al., 2014). The emergence of Vi-polysaccharide negative strains (Haque et. al., 2018) are likely to make these polysaccharide-based vaccines ineffective. Therefore, there is an urgent need to develop a highly effective typhoid vaccine, such as a vaccine containing epitopes of multiple surface proteins.

According to the Centre for Disease Control and Prevention (CDC), the development of a vaccine involves six distinct stages, such as: exploratory, preclinical, clinical development, regulatory review, approval, manufacturing, and finally, quality control (CDC, 2020).

Before going to the manufacturing level of a vaccine, it should be ensured that the safety and efficacy of that vaccine were strictly considered. The multi epitope vaccines that are designed in silico is much safer, because various computational tools are used to ensure that the putative vaccine structure is free from any harmful contents, such as allergen and toxin. By using in silico tools, codon optimization can be performed to ensure high expression of epitope vaccine. The production of this vaccine can then be easily upscaled using recombinant DNA technology. The vaccines that are developed by combining the power of bioinformatic and recombinant DNA technology are more effective, safer and cheaper than vaccines that are produced using conventional techniques (Li et. al., 2010).

Contrary to conventionally produced subunit vaccines, the localization of the in silico designed multi epitope vaccine can be specified precisely by using different bioinformatic tool. This is done to ensure that the epitopes are accessible to the components of immune system, so that upon inject of these epitopes, immune reactions can be elicited. The epitopes showing good antigenicity scores during in silico study can be ineffective if the antibodies and the immune cells can not reach them when the pathogen invasion occurs in the human body. Since a multi epitope based peptide vaccine can activate different components of immune system, it can provoke multi-dimensional attack against a pathogen. Another great advantage of such vaccine is that the pathogen has very little chance to develop resistance against it, because the pathogen will have to mutate multiple genes of it, and most mutations are deadly for the pathogen.

However, the peptides are much smaller in size compared to a protein, contributing to their lower capacity to induce immune response. For this reason, adjuvants are joined to the vaccine construct so that it can persist in human body for longer, as well as can elicit immune response more strongly. Validating the vaccine by means of molecular docking, molecular dynamics and immune simulation is very useful to make decision whether to proceed with the vaccine to further stages or not.

## Materials & Methods

### Selection of Proteins for Designing Vaccine

#### Whole Genome Sequence Retrieval

In order to search for the complete genome of *S*. Typhi, NCBI (National Center for Biotechnology Information) database was chosen (https://www.ncbi.nlm.nih.gov/). NCBI provides various useful online databases for biological sequences, such as the GenBank® nucleic acid sequence database. The search for this study returned with 2257 genomes of this bacterium. From this search result, 10 (ten) whole genomes were retrieved randomly for further study. The list of these 10 genomes is given in supplementary table 1:

#### Conversion of Whole Genome into Whole Proteome and Annotation of the Whole Proteome

The 10 whole genome sequences chosen in the previous step were needed to be converted into whole proteomes, followed by proper annotation of their proteins. For this purpose, RAST (Rapid Annotation using Subsystem Technology) server was used (www.rast.nmpdr.org). RAST is a fully-automated server for annotating genomes. At first, rapid propagation and quality check was performed. No quality revision was necessary for any entry. Afterwards, similarity computation was done, followed by bidirectional best hit computation, auto assignment and computation of pairs of close homologs. The subsystem statistics of the whole proteomes suggest that there are slight changes in the proteome compositions. It implies that in order to choose proteins for multi epitope peptide vaccine construction, the conservancy of the amino acid sequences of the chosen proteins should be given emphasize.

#### Estimation of Protein Localization using PSORTb v3.0.2

In order to be an effective epitope to be considered for constructing multi epitope vaccine, the epitope should be localized in a region of bacterial cell that is exposed to the immune system. For this purpose, estimation of protein localization and categorization of the proteins based on their position in the cell were performed using PSORTb v3.0.2 (https://www.psort.org/psortb/).

#### Assessment of Antigenicity of the Surface Proteins

The antigenicity of the surface proteins was determined using Vaxijen v2.0 (http://www.ddg-pharmfac.net/vaxijen/VaxiJen/VaxiJen.html). This tool adopts alignment-free approach for antigen prediction, which is based on auto cross covariance (ACC) transformation of protein sequences into uniform vectors of principal amino acid properties. The higher the antigenicity score, the better the chance that the proteins will provide protective antigens. In this case, the threshold for the provided dataset was set to 0.4. Proteins showing antigenicity score better than 0.4 were presumed to be probable antigens. From the antigenicity scores, top five proteins (peptidoglycan-associated lipoprotein, outer membrane protein S1, outer membrane protein F, hemagglutinin, and outer membrane porin L) having highest antigenicity score were chosen for further studies.

#### Screening of Human-similar proteins using BLASTp

If the epitopes that are chosen for multi epitope vaccine construction are similar to a protein of human protein, there is a risk that the vaccine upon entry into human body will elicit autoimmunity. For this purpose, it is important to screen out any proteins that are similar to human proteins. The BLASTp tool of NCBI was used in this case. (https://blast.ncbi.nlm.nih.gov/Blast.cgi?PAGE=Proteins). “Reference proteins (refseq protein)” database was used for protein-protein BLAST. The expected threshold value (E) was kept default as 10. Other scoring parameters such as matrix, gap costs and compositional adjustments were set as BLOSUM62, Existence:11 Extension: 1 and conditional compositional score matrix, respectively.

### Selection of Suitable Epitopes

#### Prediction of IFN- γ Epitopes

IFN-γ (Interferon Gamma) is the most important cytokine that induces both innate and adaptive immunity. They are involved in eliciting anti-tumor, antiviral and immune regulatory activities. For this purpose, identification of potential IFN-γ inducing epitopes is crucial for designing an effective subunit vaccine. Thus, the IFN-γ inducing epitopes in the target proteins were predicted using IFNepitope server (http://crdd.osdd.net/raghava/ifnepitope/). In this study, the length/window of overlapping peptides/epitopes was set as 15. Motif and SVM hybrid-based approach was adopted. IFN-gamma versus Non IFN-gamma model was chosen for the prediction.

#### CTL Epitopes Prediction

Too small epitopes will not elicit enough immune response, while too large epitopes will lead to immune tolerance. The peptide lengths for MHC-1 associated epitopes are 9-15 amino acids long, while for MHC-II, it can vary up to 22 amino acids (Arya et. al.. 2020). The NetCTL 1.2 server predicts CTL epitopes in submitted protein sequences (http://www.cbs.dtu.dk/services/NetCTL/). This server integrates prediction of peptide MHC class I binding, proteasomal C terminal cleavage and TAP transport efficiency. The server allows for predictions of CTL epitopes restricted to 12 MHC class I supertype, such as: A1, A2, A3, A24, A26, B7, B8, B27, B39, B44, B58 and B62. The 9-mer CTL (Cytotoxic T Lymphocyte) epitopes that are recognized by the most common HLA (Human Leukocyte Antigen) Class I supertypes in human population were predicted for the chosen proteins in this study. The thresholds for different parameters such as Transporter Associated with Antigen Processing (TAP) transport efficiency, proteasomal C-terminal cleavage, and epitope identification were set to 0.05, 0.15 and 0.75 respectively. In addition to this, the epitopes recognized by other HLA Class I alleles such as A-02:01, B-35:01 B-51:01 and B-58:01 were also identified utilizing Immune epitope database (IEDB) consensus method (http://tools.iedb.org/mhci). The peptides showing the consensus score of ≤ 2 were considered as strong binders and were selected for further analysis.

#### HTL Epitopes Prediction

The NetMHCIIpan 3.2 server predicts binding of peptides to MHC class II molecules (http://www.cbs.dtu.dk/services/NetMHCIIpan-3.2/). In this server, prediction services are available for the three human MHC class II isotypes HLA-DR, HLA-DP and HLA-DQ, as well as mouse molecules (H-2). The epitopes of 15-mer length recognized by HLA Class II DRB1 alleles: 01:01, 03:01, 04:01, 07:01, 08:03, 10:01, 11:01, 12:01, 13:02, 14:01 and 15:01 were predicted in this study. Besides that, the IEDB consensus method was also applied. This method employs different approaches to predict MHC Class II epitopes, including a consensus approach which combines NN-align (Neural Network-align), SMM-align (Stabilization Matrix Method-align) and Combinatorial library methods. The peptides were categorized as strong, intermediate and non-binders based on the basis of percentile rank. The threshold for strong, weak and non-binding epitopes was kept 2%, 10% and greater than 10%, respectively.

#### Selection and Merging of Promiscuous & Overlapping CTL-HTL Epitopes

It would be meaningless if the final vaccine construct is a bulky one with unnecessary and redundant sequences. It will not only be less efficient, but also costly. Hence, it would be the best if the vaccine construct is economical in size, as well as it contains as many CTL and HTL epitope regions as possible. In order to make this happen, it is necessary to select those epitopes that contain both CTL and HTL inducing regions in them. For this purpose, the starting and ending positions of the CTLs and HTLs in each protein were manually assessed and merged accordingly. These merged single peptide fragments will now be able to bind to more MHC Class I and Class II alleles and supertypes. From this approach, total 41 epitopes (5 from Peptidoglycan-associated lipoprotein, 15 from Outer membrane protein S1, 10 from Outer membrane protein F, 5 from Hemagglutinin and 7 from Outer membrane porin L) were selected for further investigation.

#### Assessment of the Antigenicity of the Epitopes

The antigenicity of 41 merged epitopes selected in the previous step was determined using Vaxijen v2.0. The antigenicity prediction threshold was kept to 0.4.

#### Peptides conservancy analysis of the T cell Epitopes

The Immune Epitope Database (IEDB) Conservancy analysis tool was used for the identification of the degree of conservancy of the predicted CTL and HTL epitopes (http://tools.iedb.org/conservancy). Here epitope linear sequence conservancy was used as analysis type. The sequence identity threshold was kept as >= 100.

#### Assessment of Population Coverage of the T cell Epitopes

The population coverage of the selected epitopes based on HLA genotypic frequencies present commonly in the world population was calculated using IEDB population coverage (http://tools.iedb.org/population). In this tool, HLA allele genotypic frequencies were obtained from Allele Frequency database (http://www.allelefrequencies.net/). Allele Frequency database provides allele frequencies for 115 countries and 21 different ethnicities grouped into 16 different geographical areas. This tool provides three calculation options for different coverage modes, such as: (1) class I separate, (2) class II separate, and (3) class I and class II combined. For each population coverage, the tool can compute the following: (1) projected population coverage, (2) average number of epitope hits/HLA combinations recognized by the population, and (3) minimum number of epitope hits/HLA combinations recognized by 90% of the population (PC90). In this study, “area_country_ethnicity” was set as query method. Class I and II was chosen for calculation option. “World” was selected as the area for population.

#### Analysis of allergenicity using AllergenFP v.1.0

In silico analysis for the prediction of the possibility of an epitope being allergenic is an important step in dry lab vaccine construction. The allergenicity of the target epitopes were evaluated using AllergenFP v.1.0 (https://ddg-pharmfac.net/AllergenFP/). In this tool, a target protein is classified as allergen or non-allergen based on the proteins with the highest Tanimoto coefficient. Since all the selected epitopes of the Hemagglutinin protein were predicted to be allergens in this tool, these epitopes were excluded for the rest of the study.

#### Selection of Epitopes for Final Vaccine Construction

After considering the antigenicity, promiscuity of the epitopes to bind with MHCs, overlapping nature, conservancy (across *Salmonella enterica* strains included in this study), population coverage and allergenicity of the predicted epitopes, top 7 T cell epitopes and top 4 IFN-γ epitopes were chosen for final vaccine construction. The amino acid sequences of the selected epitopes were compiled together for final vaccine construction.

### Similarity Search of Selected Epitopes among Other *Salmonella* Serovars

The chosen T cell and IFN-γ epitopes were subjected to protein BLAST against *Salmonella* Paratyphi A (taxid:54388), *Salmonella* Typhimurium (taxid:90371) and *Salmonella* enteritidis (taxid:149539). The parameters for BLASTp were kept as default.

#### Designing of Vaccine Primary Sequence

The promiscuous and overlapping T cell epitopes were joined using AAY linkers. The IFN-γ epitopes were linked together using GPGPG linkers. The T cell epitope region and the IFN-γ epitope region were merger using AAY linker. Adjuvant is a crucial part of a vaccine. Adjuvants are linked to a vaccine in order to increase the immunogenicity of the vaccine. The lipid A moiety of purified *S. Enterica* Serovar Typhi LPS (Lipopolysaccharide) is a potent TLR4 agonist (Bignold et. al., 1991). In order to mimic the similar immune reaction as of a real *S. enterica* Serovar Typhi infection, the multi epitope vaccine should contain TLR-4 agonist as adjuvant. In a study (Shanmugam et. al.., 2012), several Toll Like Receptor-4 (TLR-4) agonist peptides were identified. The peptide sequence GLQQVLL was the best TLR-4 stimulating peptide agonist according to that study. The peptide sequence GLQQVLL was used as the adjuvant of the proposed vaccine construct in this study. The adjuvant was joined to the vaccine using EAAAK linker. In order to purify the peptide vaccine from crude extract easily during in vitro production, 6 histidine molecules were attached to the terminal end of the vaccine construct. Overall, the peptide vaccine was composed of 201 amino acids.

### Prediction of B Cell Epitopes, Immunological and Physicochemical Properties

#### Prediction of Continuous B Cell Epitopes

After designing the peptide vaccine sequence, it is important to scan for the regions that can stimulate B cell lymphocytes in order to differentiate the cells into memory B cells and plasma cells. The BCPREDS server was used to predict linear B-cell epitopes (http://ailab-projects1.ist.psu.edu:8080/bcpred/predict.html). The BCPREDS server allows users to predict B-cell epitopes in a peptide by utilizing different prediction methods. FBCPred (Flexible length epitope prediction) was chosen as prediction method.

#### Prediction of Immunological Properties of the Designed Peptide Vaccine

The antigenicity of the designed multi epitope peptide vaccine was estimated using Vaxijen v2.0. The threshold for the candidate vaccine being antigenic was set to 0.4. Besides that, the allergenicity of the candidate vaccine was predicted using AllergenFP v.1.0.

#### Estimation of the Physicochemical Properties of the Candidate Vaccine

Different physicochemical properties of the vaccine such as theoretical isoelectric point (pI), in vitro and in vivo half-life, amino acid composition, molecular weight, instability and aliphatic index, and grand average of hydropathicity (GRAVY) of the designed candidate vaccine were estimated using the ProtParam tool of Expasy, Swiss Institute of Bioinformatics (https://web.expasy.org/protparam/).

### Structural Design of the Candidate Vaccine

#### Prediction of the Secondary Structure of the Peptide Vaccine

The distribution of different states (helices, sheets, turns, coils and acts) of the secondary structure for each recombinant epitopic structure was studied using the self-optimized prediction method (SOPMA) (https://npsa-prabi.ibcp.fr/cgi-bin/). It is based on a database that consists 126 chains of non-homologous proteins. Upon entry of a query primary sequence, SOPMA searches for similarity between the submitted protein sequence and the proteins stored in the database. Various parameters such as conformational states, similarity threshold and window width were kept as default.

#### Prediction of the Three-Dimensional Structure of the Peptide Vaccine

RaptorX server was used for the prediction of the 3D model of the peptide vaccine (http://raptorx.uchicago.edu/). It is a distance-based protein folding server facilitated by deep learning. Given an input sequence, RaptorX predicts its secondary and tertiary structures, contacts, solvent accessibility, disordered regions and binding sites.

#### Refinement of the Three-Dimensional Structure of the Peptide Vaccine

The refinement of the model retrieved was carried out using Galaxy Refine server (http://galaxy.seoklab.org/cgi-bin/submit.cgi). This server performs the repacking and molecular dynamics simulation to relax the structure, a CASP10 based refinement technique. “Conservative” was selected as the refinement option. Harmonic function was used for stronger restraint. Structures distant from the initial model were penalized.

#### Validation of the 3D Structure of the Peptide Vaccine

The validation of the tertiary structure of vaccine was carried out using ProSA-web (https://prosa.services.came.sbg.ac.at/prosa.php). The program generates scores reflecting the quality of protein structures. It estimates the overall quality of the model which is indicated in the form of z-score. If the z-scores of the predicted model are outside the range of the characteristic for native proteins, it indicates the erroneous structure. Further the Ramachandran plot analysis of the predicted model of the vaccine was carried out to using Maestro Version 11.8.012. This tool is useful for identifying residues that fall in disallowed regions of protein dihedrals, so that adjustments can be made to the protein geometry.

#### Prediction of Discontinuous B Cell Epitopes of the Vaccine

ElliPro tool of IEDB was used to predict discontinuous B cell epitopes (http://tools.iedb.org/ellipro/). This tool can predict B cell epitopes based on the 3D structure of a target protein. The protein structure can be submitted in PDB (Protein Data Bank) format. ElliPro assigns score to each predicted peptide, which is defined as a PI (Protrusion Index) value averaged over epitope residues. The three-dimensional shape of the protein is approximated by a number of ellipsoids, thus that the ellipsoid with PI = 0.9 would include within 90% of the protein residues with 10% of the protein residues being outside of the ellipsoid. For each residue, a PI value is defined based on the residue’s center of mass lying outside the largest possible ellipsoid; for example, all residues that are outside the 90% ellipsoid will have score of 0.9. Discontinuous epitopes are defined based on PI values and are clustered based on the distance R (in Å between residues’ centers of mass). The larger R is associated with the larger discontinues epitopes being predicted. The scoring threshold in the Ellipro tool for a predicted region of the peptide vaccine for being a B cell epitope was 0.5. The maximum distance R was fixed as 6 Å.

### Molecular Docking, Molecular Dynamics and Immune Simulation

#### Modelling of Toll Like Receptor-4 Protein

The three dimensional structure of TLR-4 protein was retrieved from RCSB PDB (https://www.rcsb.org/). The PDB code of the Crystal structure of the human TLR4-human MD-2-E.coli LPS Ra complex is 3FXI. This structure is composed of toll like receptor-4 protein and Myeloid Differentiation factor 2 (MD-2) protein. The MD-2 protein structure is removed from the .pdb file using PyMOL version 2.3.2, leaving the whole TLR-4 protein only. The TLR-4 structure file is saved as .pdb format for further analysis.

#### Molecular Docking of the Designed Vaccine with TLR4 Protein

The interaction between the TLR-4 and the multi epitope vaccine model was analyzed by performing molecular docking analysis using PIPER protein docking program (Kozakov et. al., 2006) from Schrodinger suite. Accurate prediction of peptide binding modes with TLR-4 protein requires accurate representation of both the peptide and the protein. In order to achieve this, both the peptide and TLR-4 protein were prepared using the Protein Preparation Wizard module of Glide (Sastry et. al., 2013). A crucial step in this preparation is the assignment of rotameric states for peptide residues with polar hydrogen atoms, such as Ser, Thr, Tyr, Asn, and Gln, as well as the orientations of water molecules, to optimize the hydrogen bonding network in the protein structure. In addition, tautomeric/ionization states of His residues are evaluated using the same criteria, and the protonation states of ionizable residues are predicted using the PROPKA algorithm (Bas et. al., 2008). The bond orders are assigned, and hydrogens are added. Water molecules are removed from the protein structures. Hetero states are generated using Epik, in pH range 7.0 +- 2.0. The PROPKA pH was set as 7.0. For energy minimization, heavy atoms were set to be converged to RMSD 0.30 Å. OPLS_2005 force field was assigned for the minimization task (Banks et. al., 2005). After minimization of the TLR-4 protein and the multi epitope peptide vaccine structures, these proteins were subjected to molecular docking. The number of ligand rotations to probe was set to 70000. Each rotation is translated to find the best docking score with the receptor. The 1000 top scoring rotations are then clustered based on binding site RMSD. Each cluster produces one docking structure. The value of maximum poses to return was set to 30. The output poses were refined appropriately.

#### Molecular Dynamics Simulation of TLR4-vaccine Complex

GROMACS (GROningen MAchine for Chemical Simulations) is a Linux-based program that is widely used for performing Molecular Dynamics Simulation (Abraham et. al., 2015). GROMACS version 2018.1 was used for molecular dynamics simulation of the TLR4-vaccine complex in order to determine whether the interaction is stable in the in vivo biological system. At first, the topology of the complex was generated. OPLS-AA (Optimized Potential for Liquid Simulation-All Atom) force field constrain was used to generate the topology file (William et. al., 1996).

In order to solvate the complex with water, a simple cubic box was defined as the unit cell. The TLR4-vaccine complex was centred in the box, and was placed at least 1.0 nm apart from the box edge. A generic equilibrated three-point water model, spc216 was used as the solvent to simulate the vaccine with periodic boundary conditions. 56844 molecules of water were required to be added to the cubic box in order to completely dissolve the complex.

The TLR4-vaccine complex has a net charge of -9e based on its amino acid composition as mentioned in their topology file. Therefore, it is necessary to add positive ions to the cubic box in order to neutralize it. GROMACS genion tool is used for this purpose. This tool can read through the topology file and replace water molecules with the ions required for neutralization of the complex. In this process, 9 Sodium (Na) molecules were added to the cubic box.

In order to ensure that the system has no steric clashes or inappropriate geometry, the structure was relaxed by energy minimization. After energy minimization, equilibration is essential by applying proper temperature based on kinetic energy, and pressure in order to have the system with proper density. At first the parameters for temperature-based equilibration were set up. NVT (isothermal-isochoric) equilibration strategy was adopted for this purpose. LINCS (linear constraint solver) for molecular simulations was used as constraint algorithm. Short-range electrostatic and van der Waals cut-offs were set as 1 nm. Particle Mesh Ewald (PME) was used for long-range electrostatics cubic interpolation (Darden et. al., 1993), and PME order was set as 4. Grid spacing for Fast Fourier Transform was set as 0.16 unit. Modified Berendsen thermostat was used as thermocouple (Berendsen et. al, 1984). The temperature for equilibration was set as 300 K (kelvin). The equilibration step was performed for 100 ps (Pico Second).

In case of pressure-based equilibration, NPT (isothermal-isobaric) equilibration strategy was adopted. The equilibration is a continuation of NVT equilibration; hence the parameters were kept similar to temperature-based equilibration. Parrinello-Rahman pressure coupling was used (Parrinello & Rahman 1980). 1 bar pressure was applied to the system. The equilibration was run for 100 ps.

Upon completion of the two equilibration phases, the system was well-equilibrated at the desired temperature and pressure. This equilibrated system was subjected to molecular dynamics simulation. The simulation was run for 2,500,000 steps (5000 ps = 5 ns). After simulation, trjconv was used as post-processing tool to examine any periodicity in the system. Using this tool, RMSD and RMSF of the system was analysed. All the graphs were visualized and presented using XMGRACE software (Turner, 2005)

#### Immune Simulation of TLR4-vaccine Complex

In order to mimic the in vivo immune reactions after vaccination, immune simulations were conducted using the C-ImmSim server (http://150.146.2.1/C-IMMSIM/index.php). It simulates the regimen and real-time dosage response patterns of a vaccine. This software exploits an agent-based model that uses a position-specific scoring matrix (PSSM) for immune epitope prediction and machine learning techniques for prediction of immune interactions. The minimum recommended time interval between two doses of vaccine is 4 weeks for most of the currently used vaccines (Castiglione et. al., 2012). Therefore, three injections were simulated to be applied four weeks apart. These injections were given at 1, 84, and 168 (each time step is 8 hours and time step 1 means injection at time = 0). The dosage was defined at 1 arbitrary unit, and the simulation volume was set as 20 micro litre. The simulation was run for 100 steps. The other parameters of simulation were kept default.

### Codon Adaptation and In Silico Cloning

#### Codon Optimization of the Multi Epitope Vaccine Construct

Reverse translation and codon optimization of the peptide vaccine were performed using Java Codon Adaptation Tool (JCat) server (http://www.prodoric.de/JCat). JCat server can optimize and estimate the adaptation of the codon usage, in order to express a foreign protein in a selected expression vector. The adaptation is based on CAI (codon adaptation index) values proposed by P.M. Sharp and W.H. Li (Sharp & Li 1987). The CAI-values were calculated by applying an algorithm proposed by A. Carbone (Carbone et. al., 2003). In this study, codon optimization was performed to express the final vaccine construct in the E. coli (strain K12) host, as codon usage of E. coli differs from that of *Salmonella* enterica Serovar Typhi. Besides, the Jcat tool was instructed to perform the reverse translation process by avoiding the rho-independent transcription termination, the algorithm for this purpose is based on a model from MD. Ermolaeva (Ermolaeva et. al., 2000). The tool is further instructed to avoid prokaryote ribosome binding site, and cleavage sites of EcoRI and EcoRV restriction enzymes. The JCat output includes the codon adaptation index (CAI). and percentage GC content. The values of these parameters can be used to assess protein expression levels.

#### In Silico Cloning of the Codon Optimized DNA

The codon optimized sequence was inserted into the pET-30a (+) vector using SnapGene tool version 5.2.4 to ensure vaccine expression. The DNA fragment was inserted in between the cleavage sites of EcoRI and EcoRV on the pET-30a (+) vector.

## Result

### Protein Targets

After retrieval of whole genomes from NCBI database, the RAST annotation server was utilized to convert these whole genomes into whole proteomes with full annotation. The proteins were sorted using PSORTb v3.0.2, and proteins that were exposed in the outer environment was preferred as target proteins. The antigenicity of the proteins was also assessed using Vaxijen server, and protein conservancy was analysed using IEDB conservancy analysis tool. Based on localization, antigenicity and conservancy, five most potent surface proteins were chosen. The above-mentioned characteristics of these proteins are mentioned in table 1.

**Table 1:** List of Protein Targets mentioning their localization, antigenicity and conservancy scores

| Protein Name | Localization in Cell | Localization Score (Out of 10) | Antigenicity (threshold = 0.40 or above) | Conservancy (Across all proteomes) |
| --- | --- | --- | --- | --- |
| Peptidoglycan-associated lipoprotein | Extracellular (Surface Protein) | 10.0 | 0.9175 | 100% |
| Outer Membrane Protein S<sub>1</sub> | Extracellular (Surface Protein) | 10.0 | 0.8837 | 100% |
| Outer Membrane Protein F | Extracellular (Surface Protein) | 10.0 | 0.8294 | 100% |
| Hemagglutinin | Extracellular (Surface Protein) | 9.95 | 0.794 | 100% |
| Outer Membrane Porin L | Extracellular (Surface Protein) | 9.93 | 0.7906 | 100% |

Besides, all these potent target proteins were subjected to BLASTp. After performing protein BLAST against human proteome, all the proteins showed 0% similarity with any single protein of *Homo sapiens*.

### T cell & IFN- γ Epitopes Prediction

#### T Cell Epitopes

The NetCTL 1.2 and IEDB consensus method were utilized for the prediction of CTL epitopes. Besides that, the HTL epitopes were predicted using Net MHC II pan 3.2 server and IEDB consensus method. After that, the epitopes which showed better binding scores were chosen for further analysis, such as conservancy, population coverage, immunogenicity and allergenicity. Based on all these parameters, the study led to the identification of some epitopes, which are enlisted in supplementary table 2. From these epitopes, the promiscuous and overlapping CTL and HTL epitopes were merged together in order to make them single fragments for vaccine design. The epitopes for the multi epitope vaccine construct were chosen based on the following criteria: (a) The epitopes should be promiscuous, (b) should have overlapping CTL and HTL epitopes, (c) antigenicity, (d) population coverage, (e) should not match with any human gene (means it should not invoke autoimmunity). On the basis of these parameters, 7 T cells epitopes (Merged peptides comprising of 7 HTL and 7 CTL epitopes, Table: 2) was selected for construction of the vaccine.

**Table 2:**
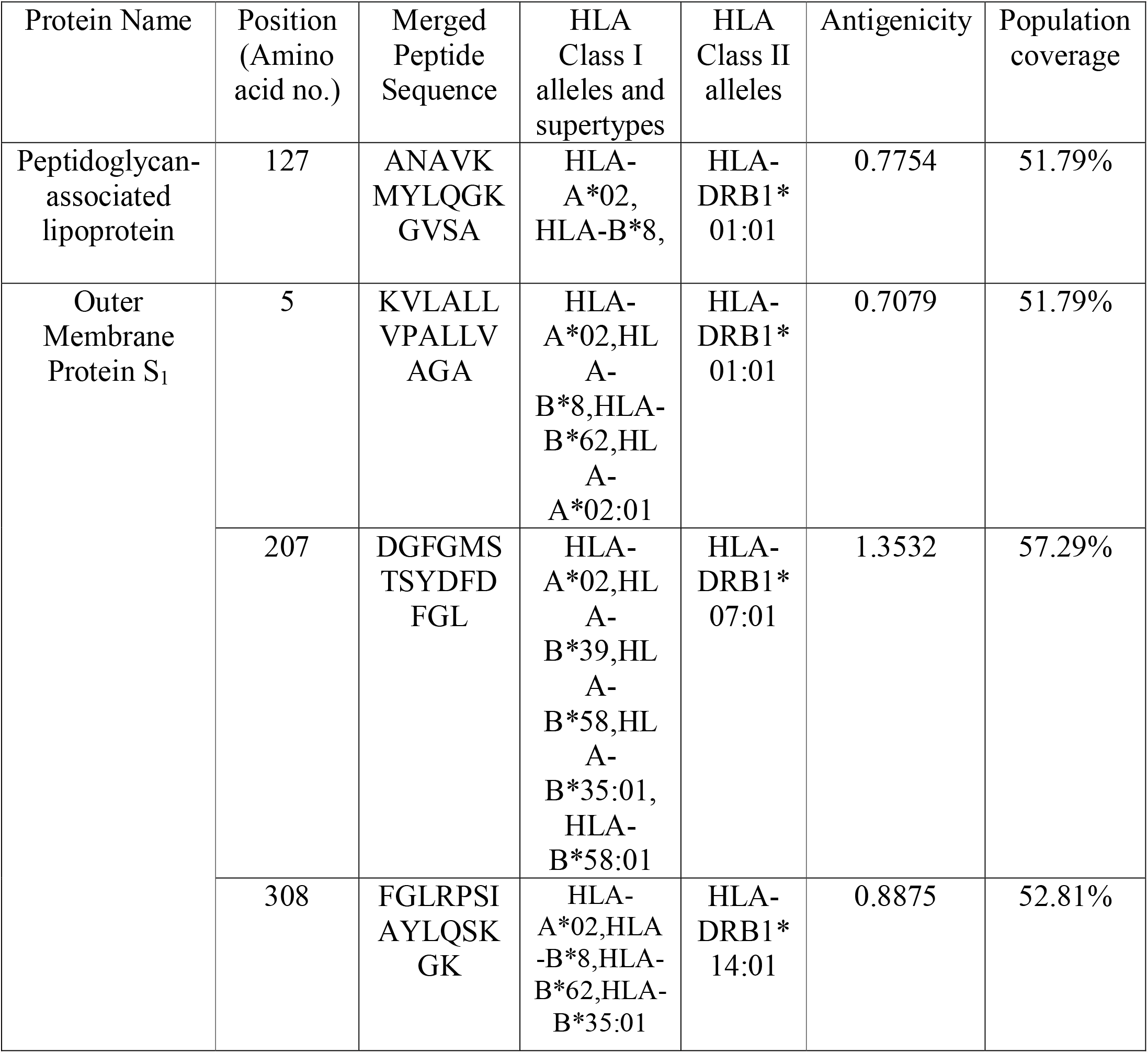
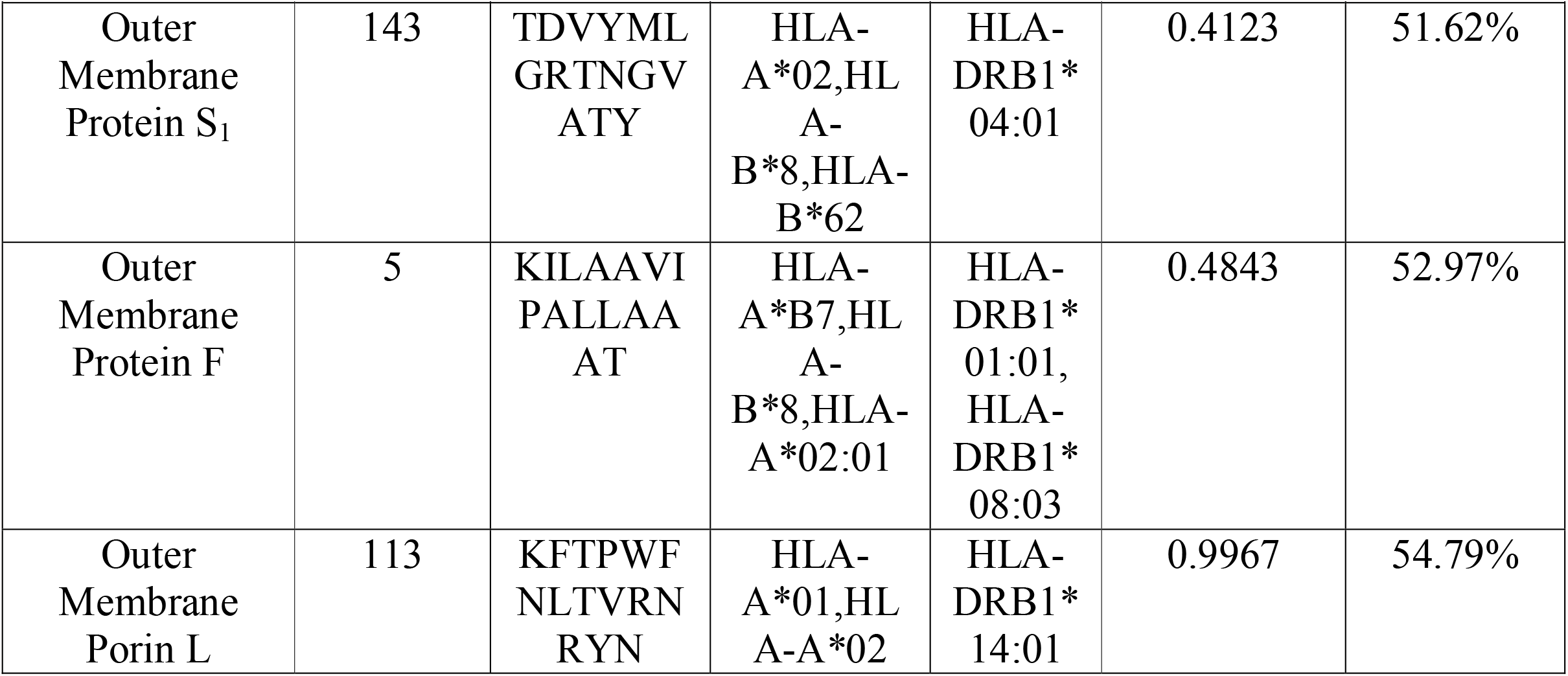
List of selected T cell epitopes along with their position, amino acid sequence, interaction with HLA Class I and II alleles, antigenicity and population coverage

The mapping of the T cell epitopes on individual proteins are illustrated in Figure 1.

**Figure 1:**
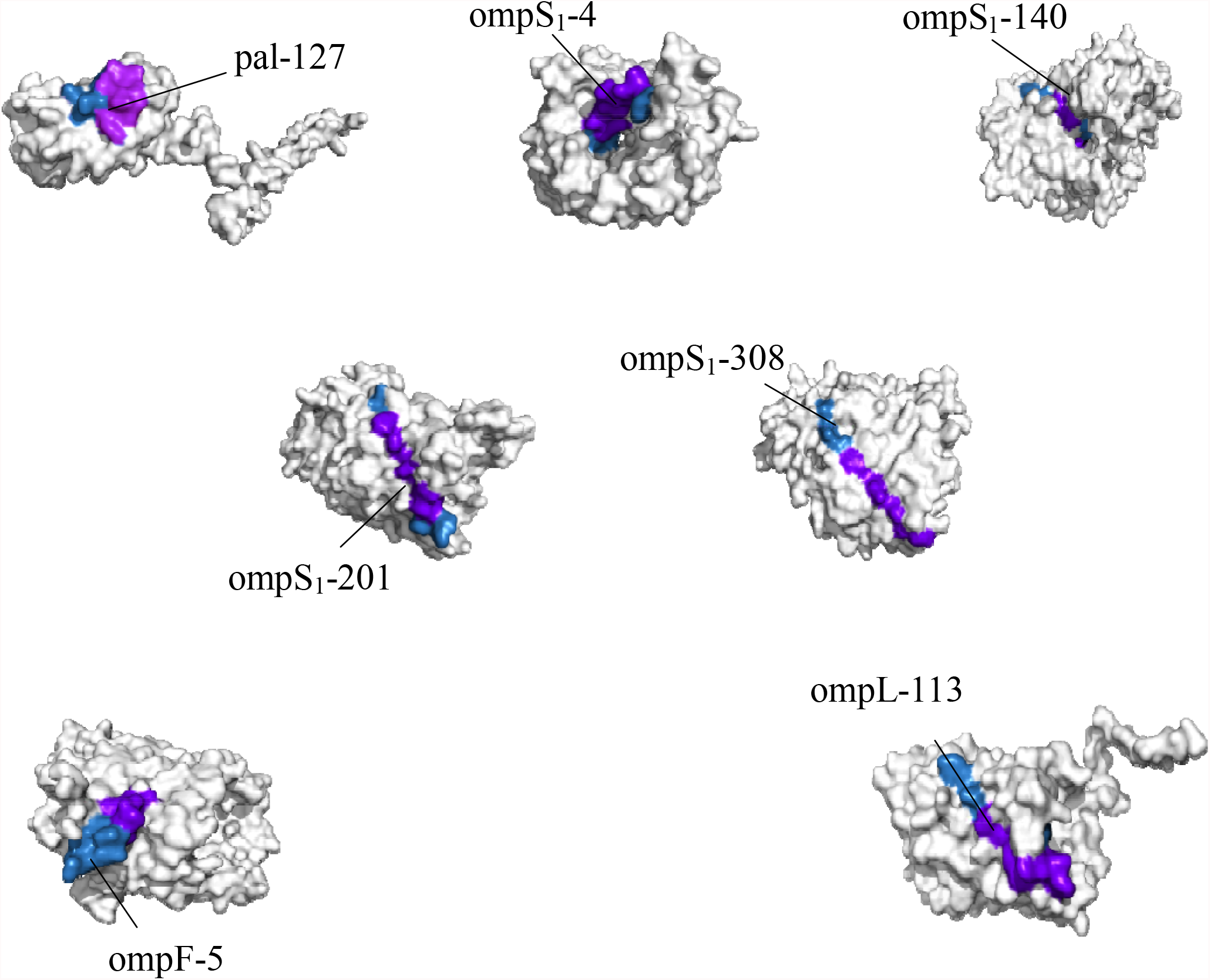
Mapping of T Cell Epitopes on Respective Proteins. Here, the full form of the abbreviations in the figure are: pal = Peptidoglycan-associated lipoprotein, ompS1 = Outer Membrane Protein S1, ompF = Outer Membrane Protein F, ompL = Outer Membrane Porin L. The number next to the abbreviations of proteins denotes the amino acid position of the starting of the epitopes in their respective proteins. In this figure, the purple + royal blue colour denotes the whole HTL epitope region, while the purple colour in the HTL epitope region denotes the 4 IFN-γ inducing epitopes overlapping CTL epitope region.

#### IFN- γ Epitopes

In addition, the IFNepitope server was employed for the prediction of IFN-γ inducing epitopes. The list of potential IFN- γ epitopes of the protein targets included in this study are mentioned in supplementary table 3. The IFN- γ epitopes included in the multi epitope vaccine are enlisted in Table 3.

**Table 3:**
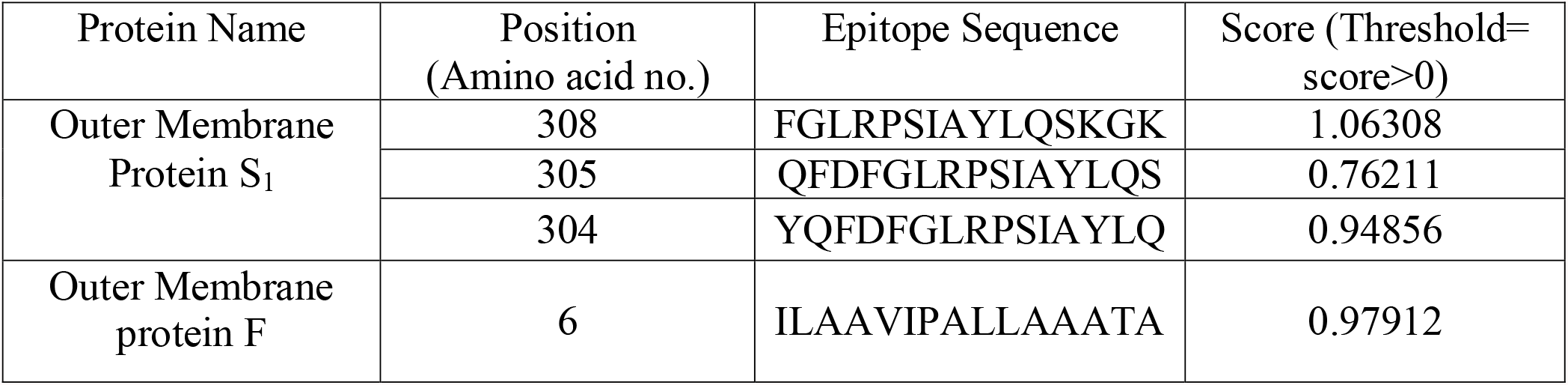
List of Potential IFN- γ Epitopes along with their potency score. The epitope sequences are listed with decreasing order of their score.

The mapping of the IFN-γ inducing epitopes on individual proteins are illustrated in Figure 2.

**Figure 2:**
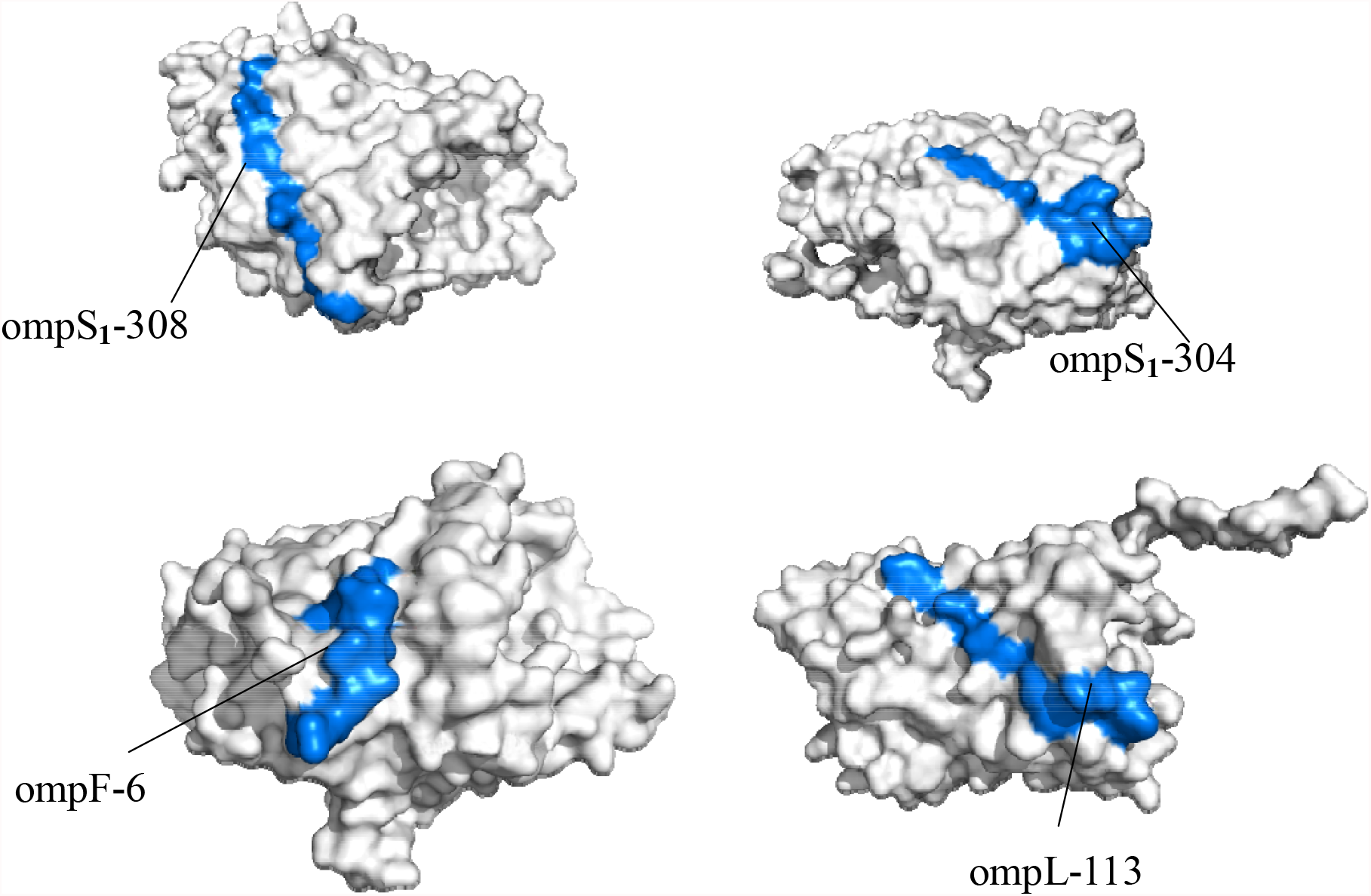
Mapping of the 4 IFN-γ inducing epitopes on Respective Proteins. Here, the full form of the abbreviations in the figure are: pal = Peptidoglycan-associated lipoprotein, ompS_1_ = Outer Membrane Protein S_1_, ompF = Outer Membrane Protein F, ompL = Outer Membrane Porin L. The number next to the abbreviations of proteins denotes the amino acid position of the starting of the epitopes in their respective proteins. In this figure, the royal blue colour denotes the 4 IFN-γ inducing epitopes

**Figure 3:**
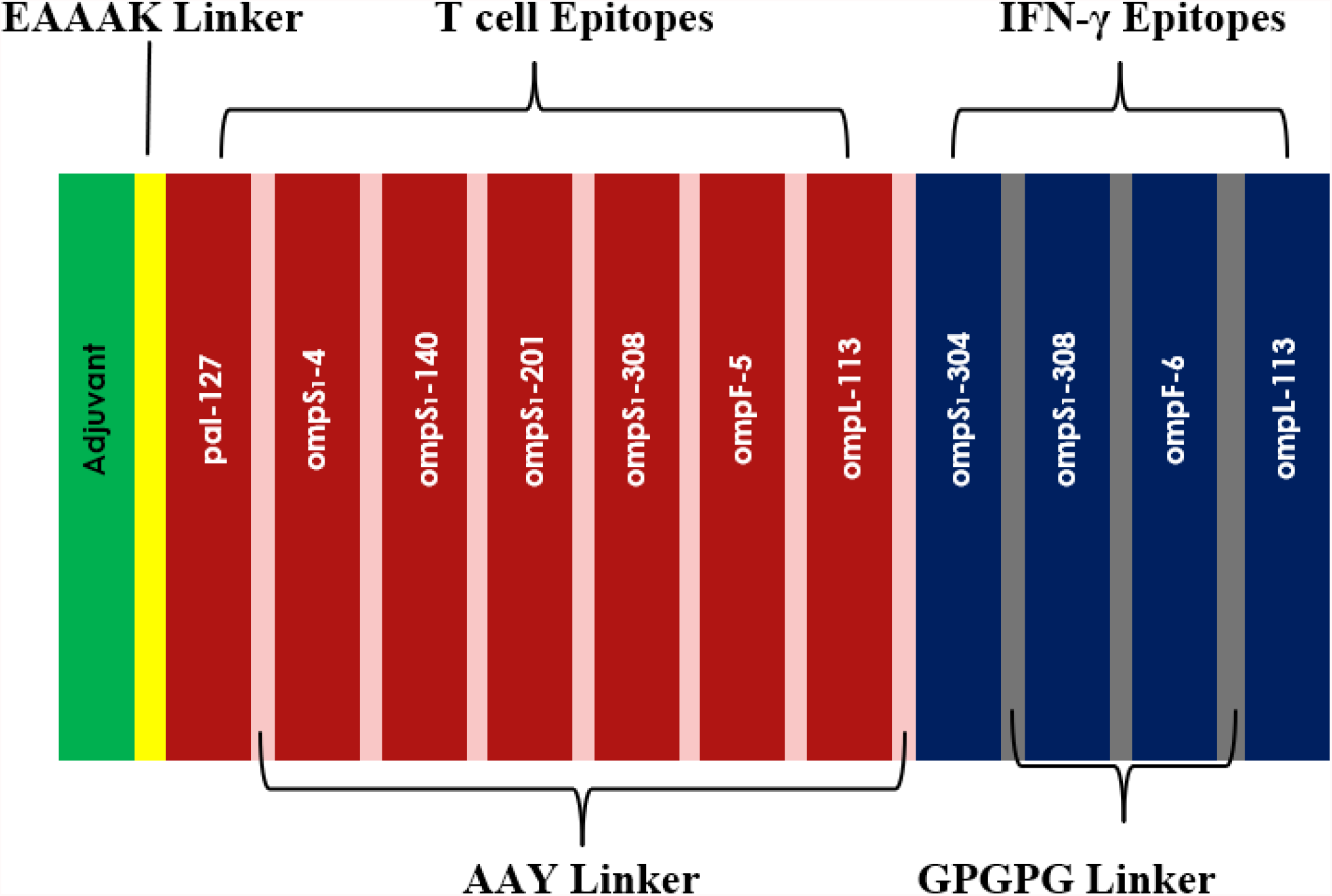
Schematic View of Final Vaccine Construct

### BLASTp of Selected T cell & IFN- γ Epitopes

BLASTp against three pathogenic serovars of *Salmonella*, such as: *Salmonella* Paratyphi A, *Salmonella* Typhimurium and *Salmonella* Enteritidis revealed that the amino acid sequence of all the selected seven T cell & four IFN- γ Epitopes of *Salmonella* Typhi are 100% similar to the epitopes of corresponding *Salmonella* serovar. This indicates that the proposed vaccine might be effective not only against *Salmonella* Typhi, but also against most common pathogenic serovars of *Salmonella*. The detailed results are mentioned in the supplementary table 4.

#### Multi Epitope Vaccine Construction, Structural Properties and Refinement

##### Prediction of Primary Sequence of the Vaccine

The T cell epitopes, IFN-γ epitopes and the adjuvant GLQQVLL peptide were joined together using suitable linkers, resulting in a continuous primary sequence of the vaccine. A stretch of 6 histidine molecules were added to the terminal region of the vaccine. This final vaccine construct was composed of 201 amino acids.

##### Estimation of Secondary Structure of the Vaccine

The secondary structural properties of the vaccine were estimated using SOPMA server. The results showed that the vaccine candidate was composed of 41.79 % alpha helical regions, 12.43% extended strands, 7.96% beta turns and 36.82% random coil.

##### Designing of Tertiary Structure of the Vaccine

The three-dimensional structure was constructed using Raptor-X online web server. The 3D structure of the vaccine is illustrated in Figure 4.

**Figure 4.**
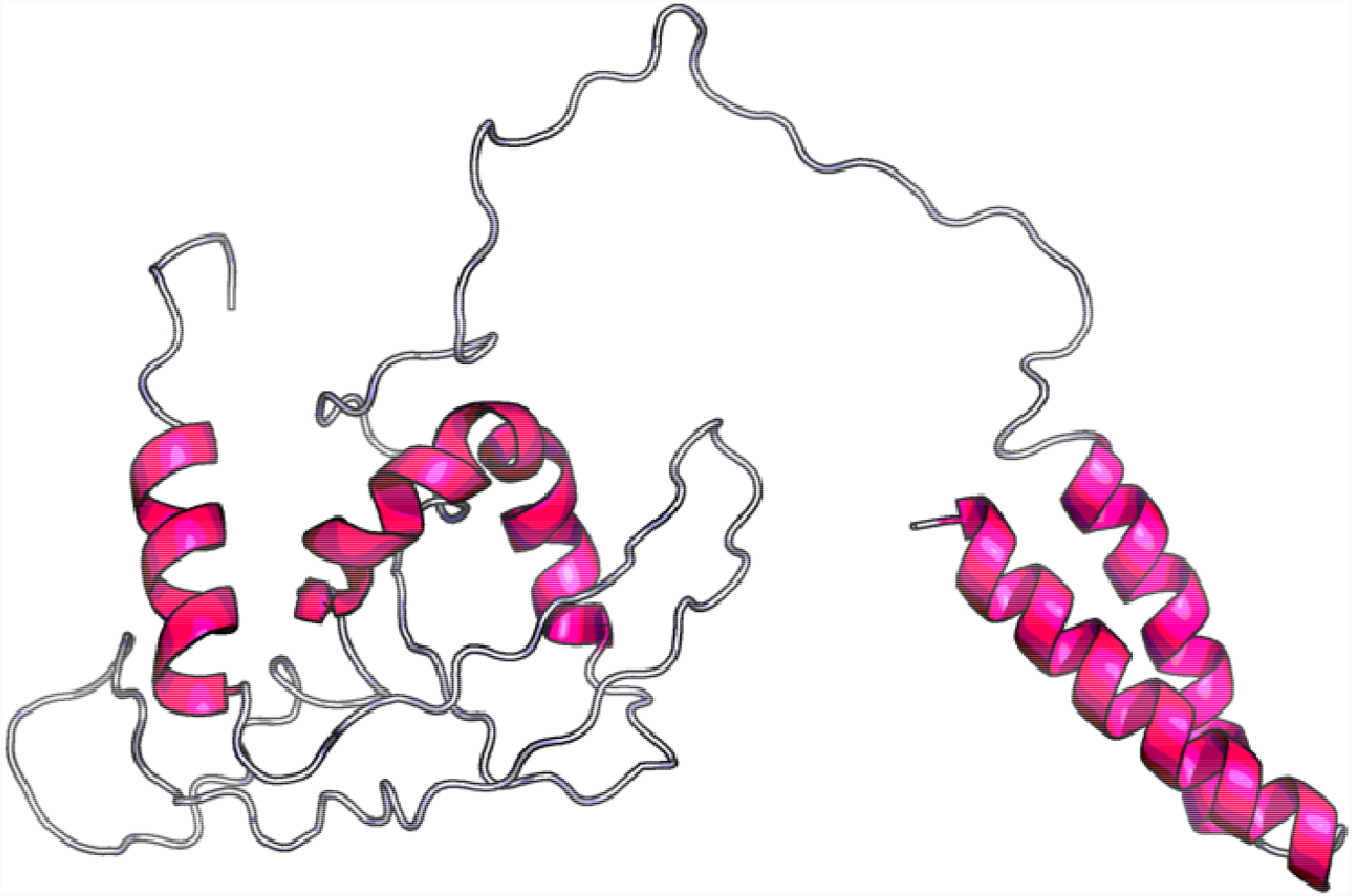
Three Dimensional Structure of the Protein Derived from RaptorX

##### Refinement of the 3D Structure

For refinement purpose, the 3D model was subjected to Galaxy Refine server 2. The server yielded 10 probable models for the vaccine, as presented in the supplementary table 5. Among the refined models, the model 6 showed the best results in terms of the overall quality of the model prepared. The 3D structure of the refined model 6 is illustrated in Figure 5.

**Figure 5:**
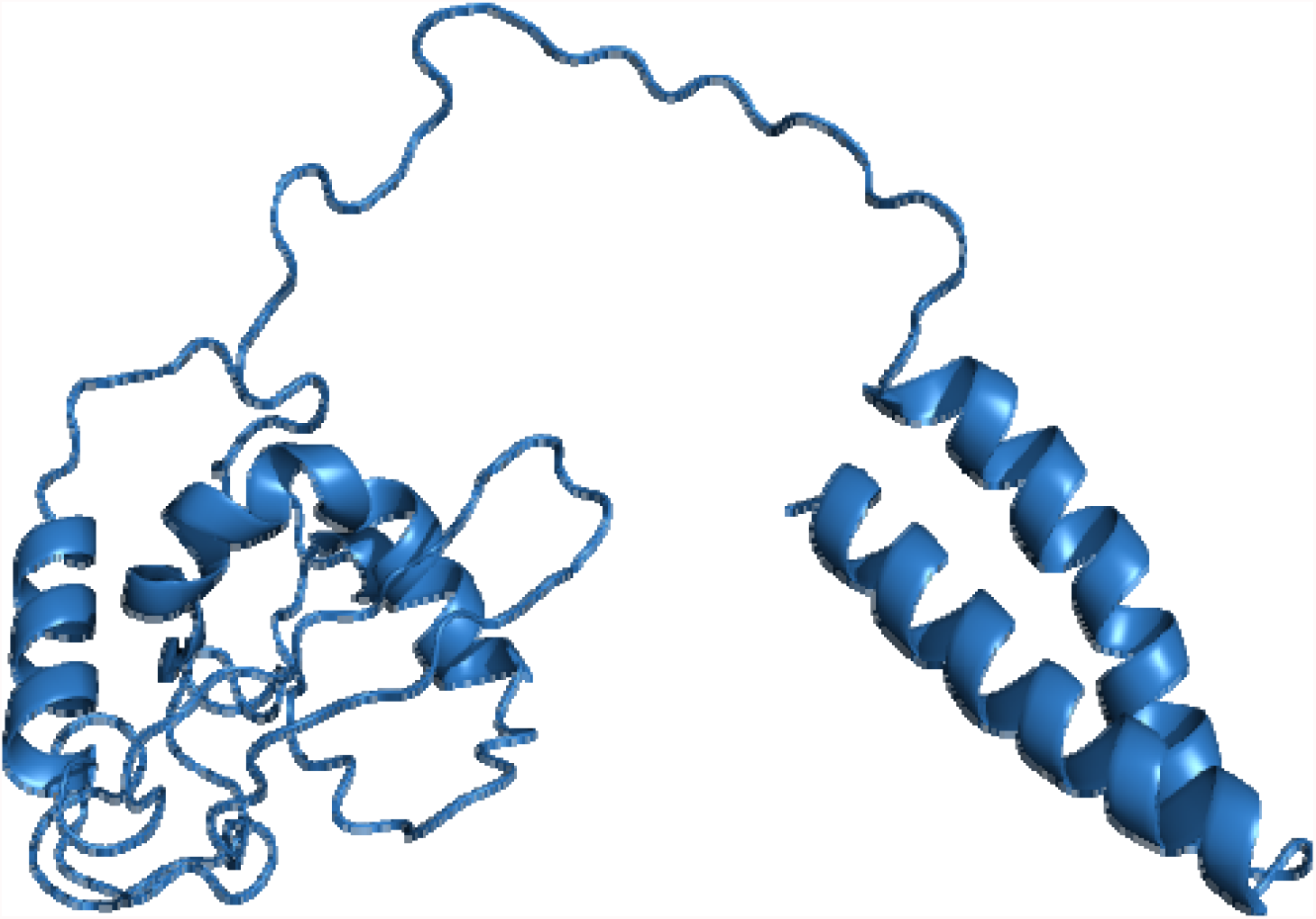
Refined Model of the Multi Epitope Vaccine Candidate

##### Validation of the 3D Structure

In order to analyze the model quality, the finalized model was subjected to ProSA-web. The local model quality graph, the energy distribution and the graph of Z score against number of residues and the Ramachandran Plot of the vaccine model are illustrated in Figure 6. The web server revealed the z score of −4.39 for the model constructed which was lying inside the scores range of the comparable sized native proteins (Figure 6c). Finally, the overall quality of the finalized model of the multi-epitope vaccine construct was analysed using Ramachandran plot analysis. The results show 96.0%, 3.5% and 0.5% residues lying in favoured, allowed and outlier regions (Figure 6d).

**Figure 6:**
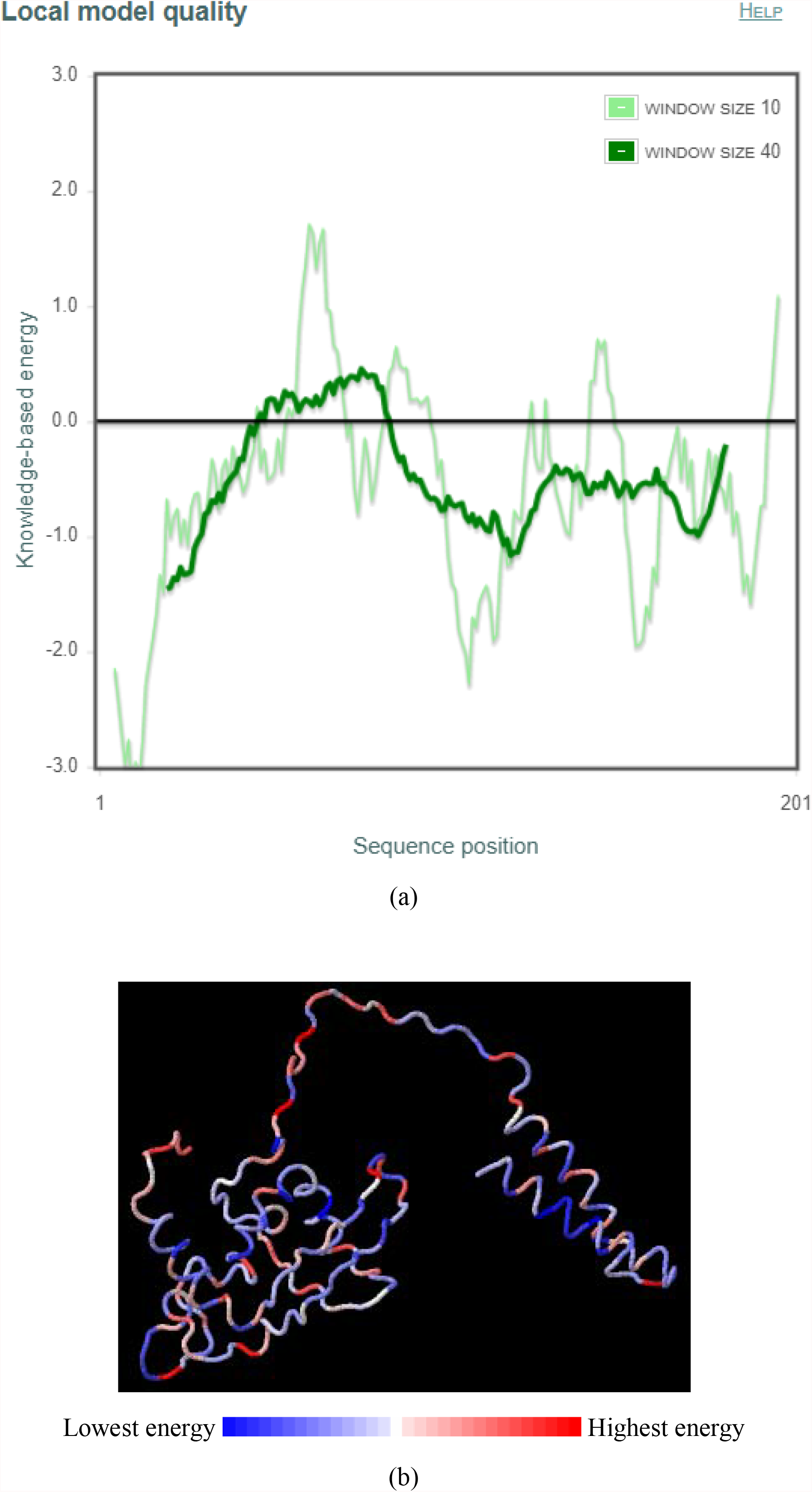
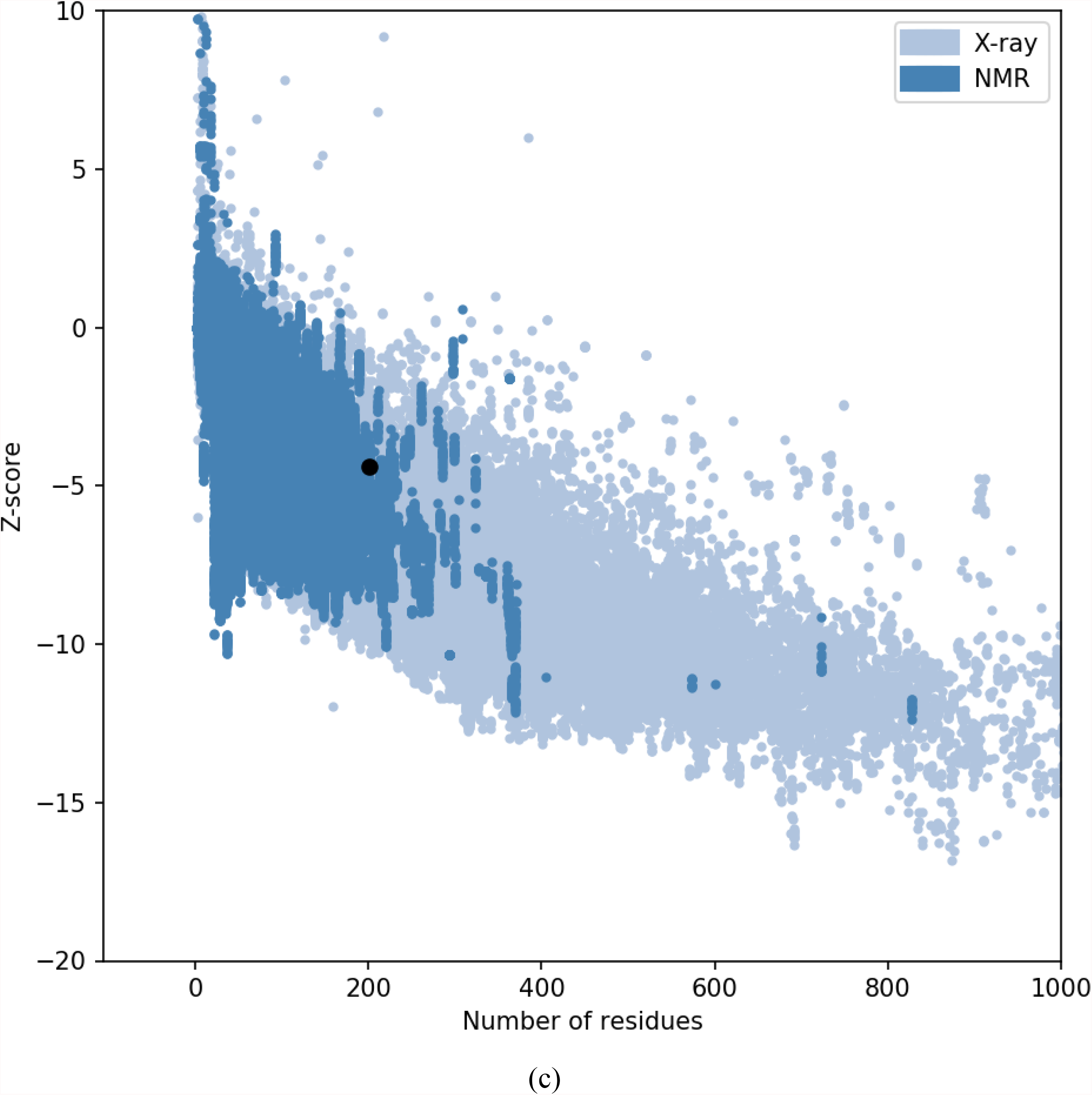
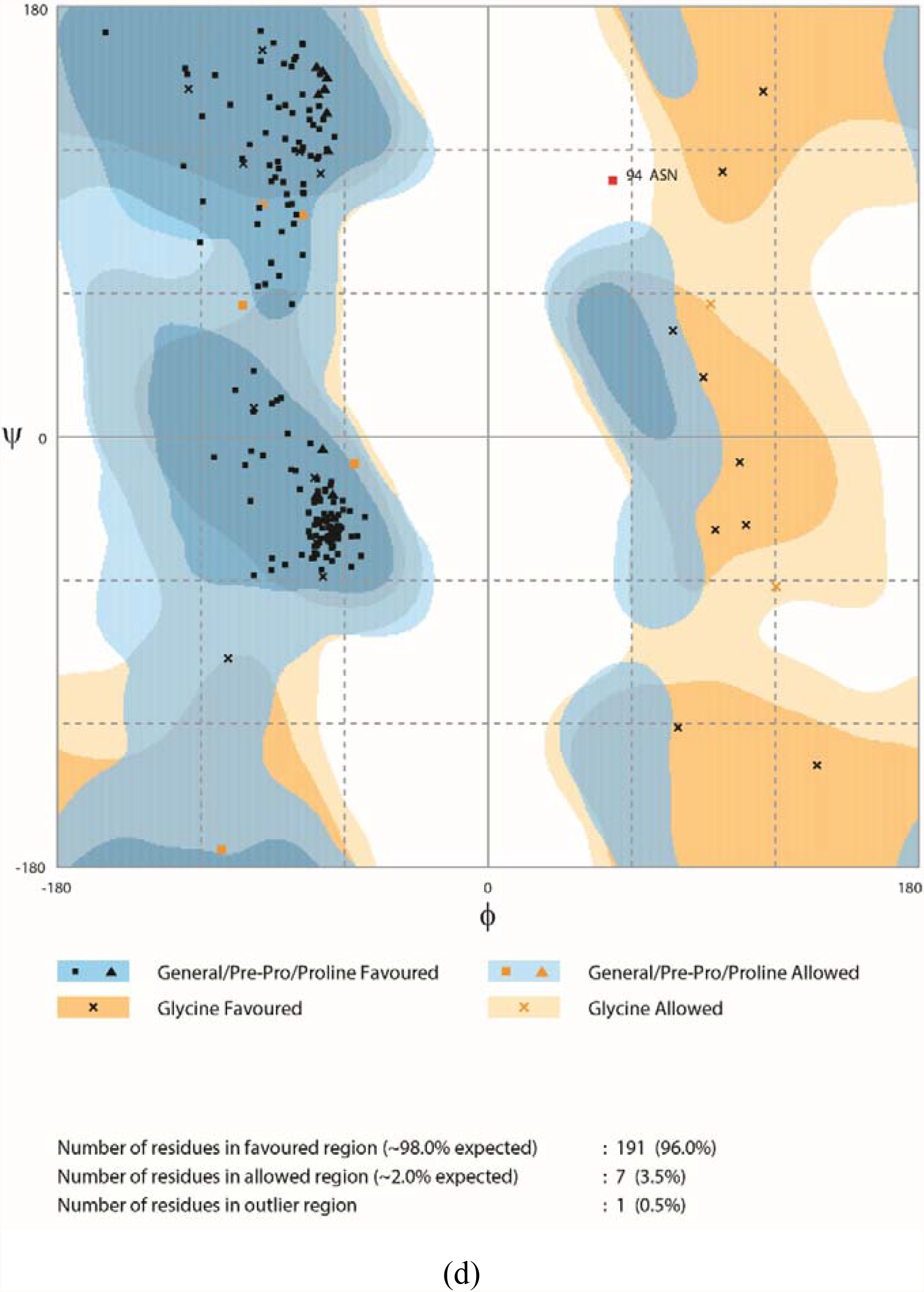
Validation of the 3D structure of the multi epitope vaccine. Here, (a) Local Model Quality Graph of the Vaccine Model, (b) Local Model Quality Graph of the Vaccine Model, (c) Validation of the structure with a Z score of −4.39 using PROSA, (d) Ramachandran Plot of the Multi Epitope Vaccine Model

#### B cell Epitope Prediction and Physicochemical properties of the final vaccine construct

##### B Cell Epitopes

The multi-epitope vaccine construct was predicted to be antigenic in the Vaxijen server, with the immunogenicity score of 0.8534. It was also predicted as non-allergen in the Allergen FP server. The linear/continuous and conformational/discontinuous B cell epitopes in the vaccine construct were predicted using BcPred 2.0 and Ellipro server. The servers predicted 4 linear/continuous B cell epitopes (Table 4) and 3 conformational/discontinuous B cell epitopes (Table 5). The mapping of the 3 conformational B cell epitopes is illustrated in Figure 7.

**Table 4:** B Cell Epitopes Derived from BcPred 2.0 in the Multi Epitope Vaccine Candidate

| Position | Epitope | Score |
| --- | --- | --- |
| 132 | NRYNGPGPGFGLRP | 1 |
| 171 | IAYLQGPGPGILAA | 1 |
| 151 | QSKGKGPGPGYQFD | 1 |
| 77 | QSKGKAAYTDVYML | 0.999 |

**Table 5:** List of Discontinuous B Cell Epitopes resulting from the Analysis in Ellipro Server

| No. | Residues | Number of residues | Score |
| --- | --- | --- | --- |
| 1 | _:A9, _:A10, _:A11, _:K12, _:A13, _:N14, _:A15, _:V16, _:K17, _:M18, _:Y19, _:L20, _:Q21, _:G22, _:K23, _:G24, _:V25, _:S26, _:A27, _:A28, _:A29, _:K31, _:V32, _:L33, _:A34, _:L35, _:L36, _:V37, _:P38, _:A39, _:L40, _:L41, _:V42, _:G44, _:A45, _:A46, _:A47, _:Y48, _:D49, _:G50, _:F51, _:G52, _:M53, _:S54, _:T55, _:S56, _:Y57 | 47 | 0.778 |
| 2 | _:D60, _:F61, _:G62, _:L63, _:A64, _:A65, _:Y66, _:F67, _:G68, _:L69, _:R70, _:S72, _:I73 | 13 | 0.755 |
| 3 | _:W125, _:N127, _:L128, _:T129, _:V130, _:R131, _:N132, _:Y134, _:N135, _:G136, _:P137, _:G138, _:P139, _:G140, _:G142, _:L143, _:R144, _:P145, _:S146, _:I147, _:A148, _:Q151, _:S152, _:K153, _:G154, _:K155, _:G176, _:P177, _:G178, _:P179, _:G180, _:I181 | 32 | 0.594 |

**Figure 7:**
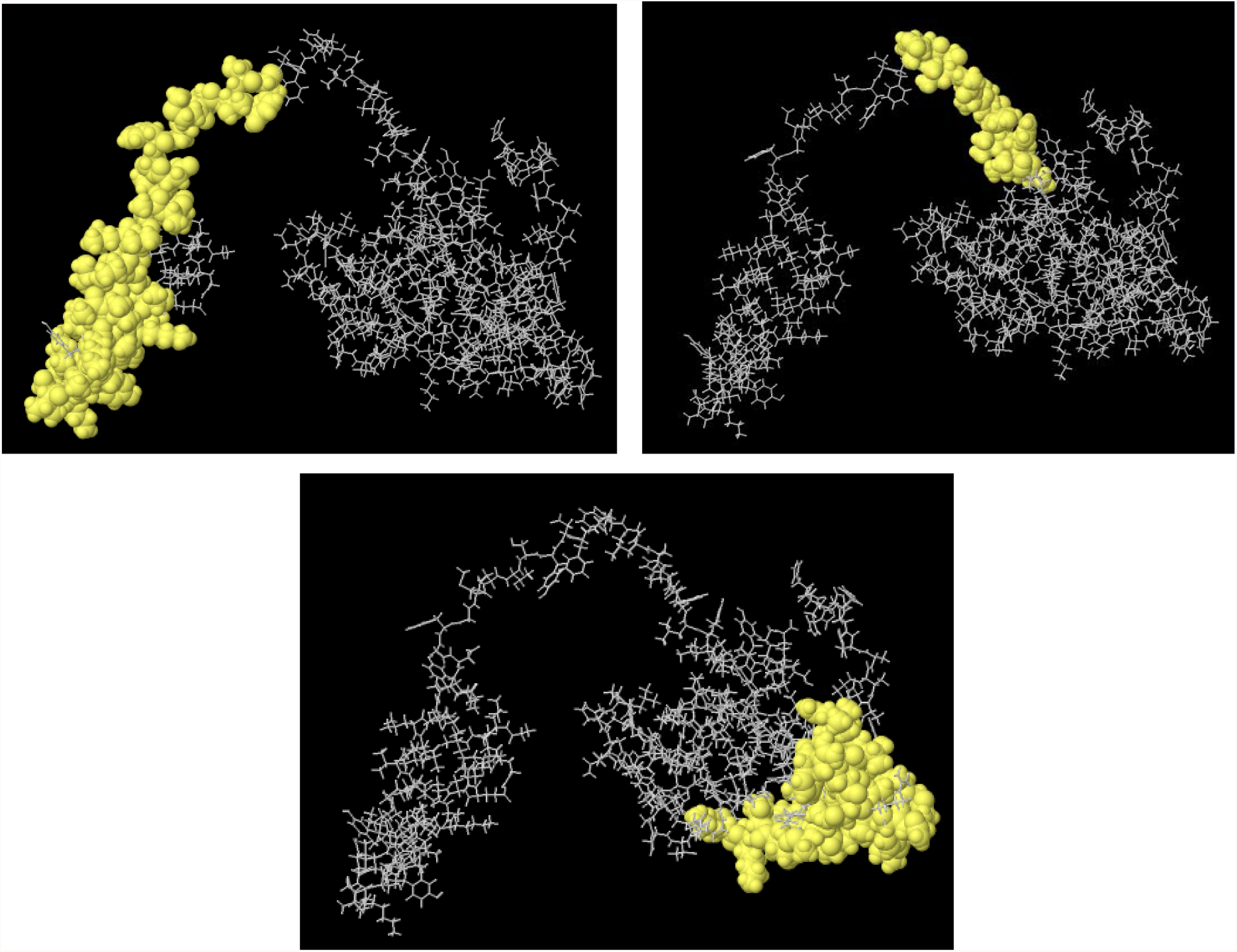
Illustration of the 3 conformational B cell epitopes on the Vaccine. The 3 conformational B cell epitope regions are marked as yellow.

#### Physicochemical Properties

The result from Protparam server for predicting physicochemical properties of the vaccine shows that the molecular weight of the vaccine candidate is 21034.39. The theoretical pI is 9.75. The total number of negatively charged residues (Asp + Glu) are 6, while the total number of positively charged residues (Arg + Lys) are 16. The chemical formula of the vaccine is C_979_H_1498_N_254_O_257_S_3._ Total number of atoms in this peptide vaccine is 2991. The extinction coefficient value was measured as 27850, and are expressed in units of M-1 cm-1, at 280 nm measured in water. The estimated half-life of the vaccine inside mammalian reticulocytes, in vitro is 30 hours. The instability index (II) is computed to be 38.96. This classifies the protein as stable. The Aliphatic index and the Grand average of hydropathicity (GRAVY) are predicted to be 95.42 and 0.296, respectively.

### TLR-4 Modelling and Molecular Docking Interaction with the Final Vaccine Construct

The Toll like Receptor-4 protein and the multi epitope vaccine model were docked together using the piper protein-protein docking module of Schrodinger Maestro. This docking tool generated 30 docking poses between TLR-4 and vaccine model. The models were ranked based on PIPER pose energy. The list of the PIPER pose energy, PIPER pose score, PIPER cluster, Potential Energy based on OPLS3e force field and RMS Derivative based on OPLS3e are mentioned in supplementary table 6. The best docking pose between the TLR-4 protein and the multi epitope vaccine construct are illustrated in Figure 8.

**Figure 8:**
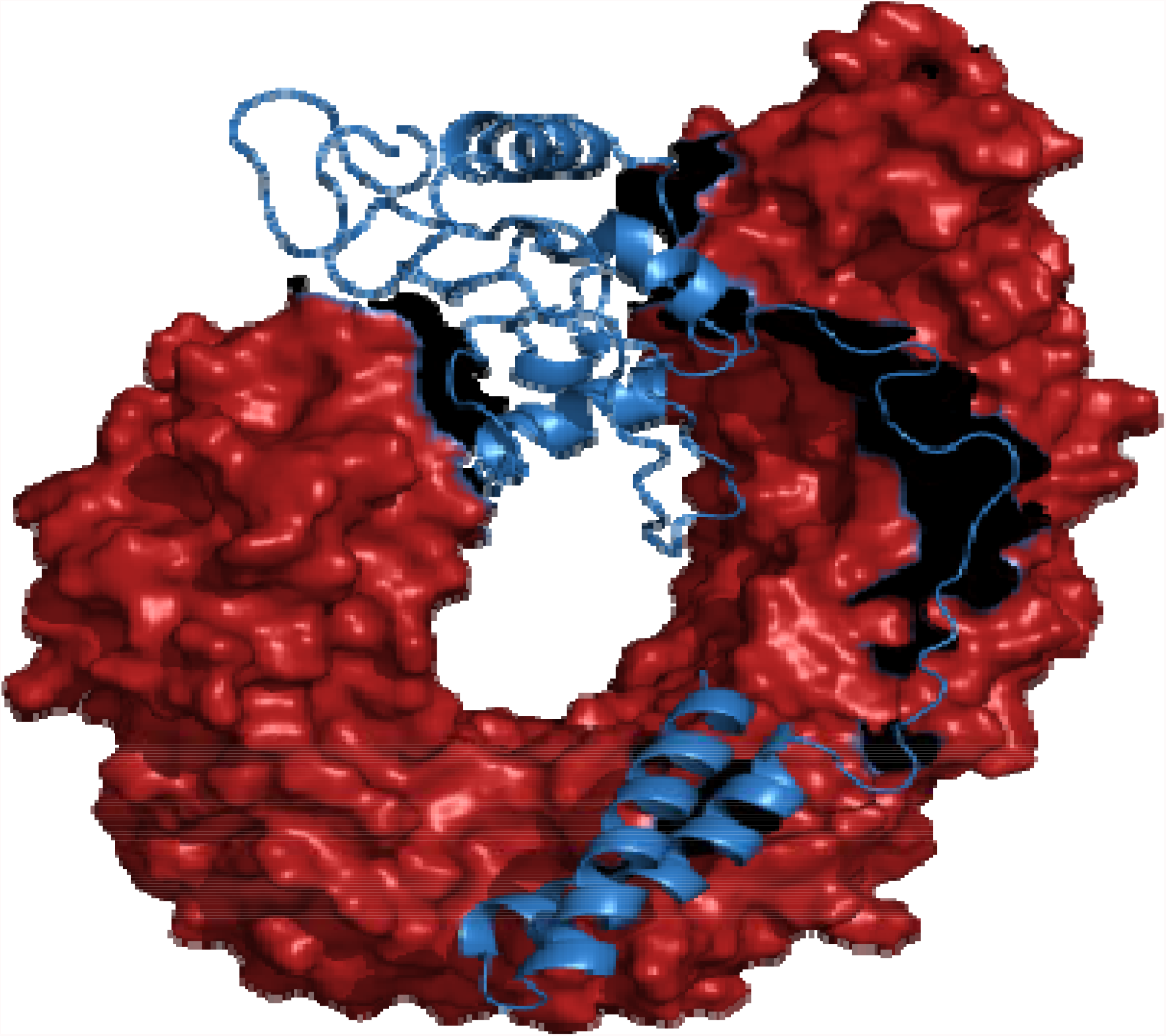
Docked structure of the TLR-4 Protein and Multi Epitope Vaccine Structure. The Red protein indicates TLR-4 protein, while the blue structure indicates vaccine peptide

The interaction types between TLR-4 and vaccine, along with associated details are enlisted in Supplementary Table 7.

### Molecular Dynamics

The molecular dynamics simulation was performed to examine the physical movements of atoms and molecules of the final multi epitope vaccine structure. The structure was energy minimized accordingly by applying proper temperature and pressure mimicking real life scenario. The stability of the structure was analysed by calculating the root square deviation (RMSD) of the protein backbone and the root mean square fluctuation (RMSF) of all side chain residues of the multi epitope vaccine construct. The RMSD of the complex of TLR-4 (receptor) and multi-epitope vaccine (ligand) was found to be ∼3.5 Å (Figure 9a), indicating that the structure is very stable. Further the RMSF of amino acid side chains was calculated to estimate the stability of the ligand-receptor interaction (Figure 9b). The RMSD plot shows that the fluctuations are minimal, indicating the uninterrupted interaction between ligand and receptor. In contrast, the fluctuations in the RMSF plot were very high indicating the flexibility in the regions between ligand-receptor complex. In RMSF plot, the initial peaks were very high (even greater than 6 Å) indicating the high degree of flexibility in the receptor-ligand complex.

**Figure 9a:**
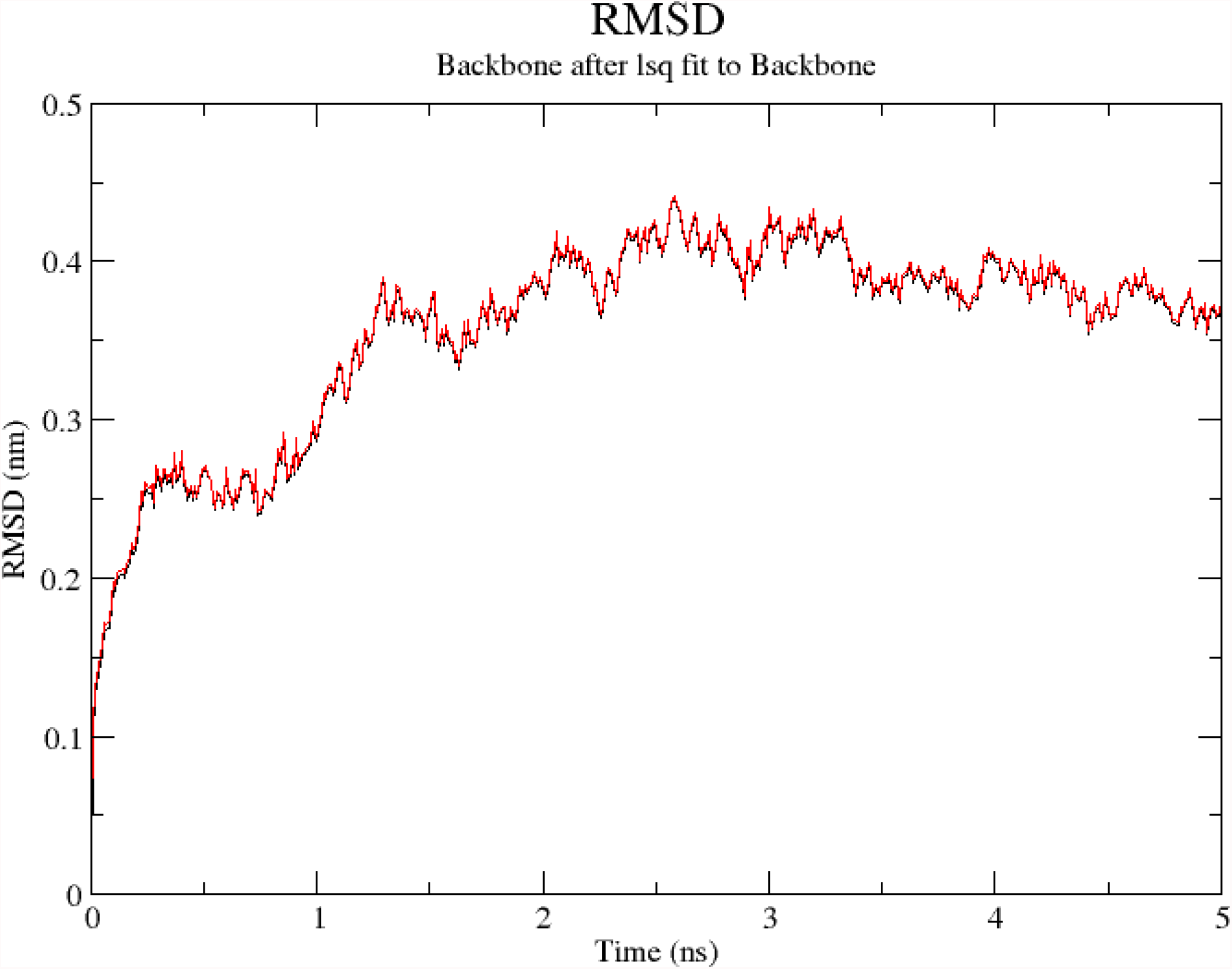
Comparative RMSD graph between the crystal structure of TLR 4-Vaccine complex (Red) and the MD simulated structure of TLR 4-Vaccine complex (Black)

**Figure 9b:**
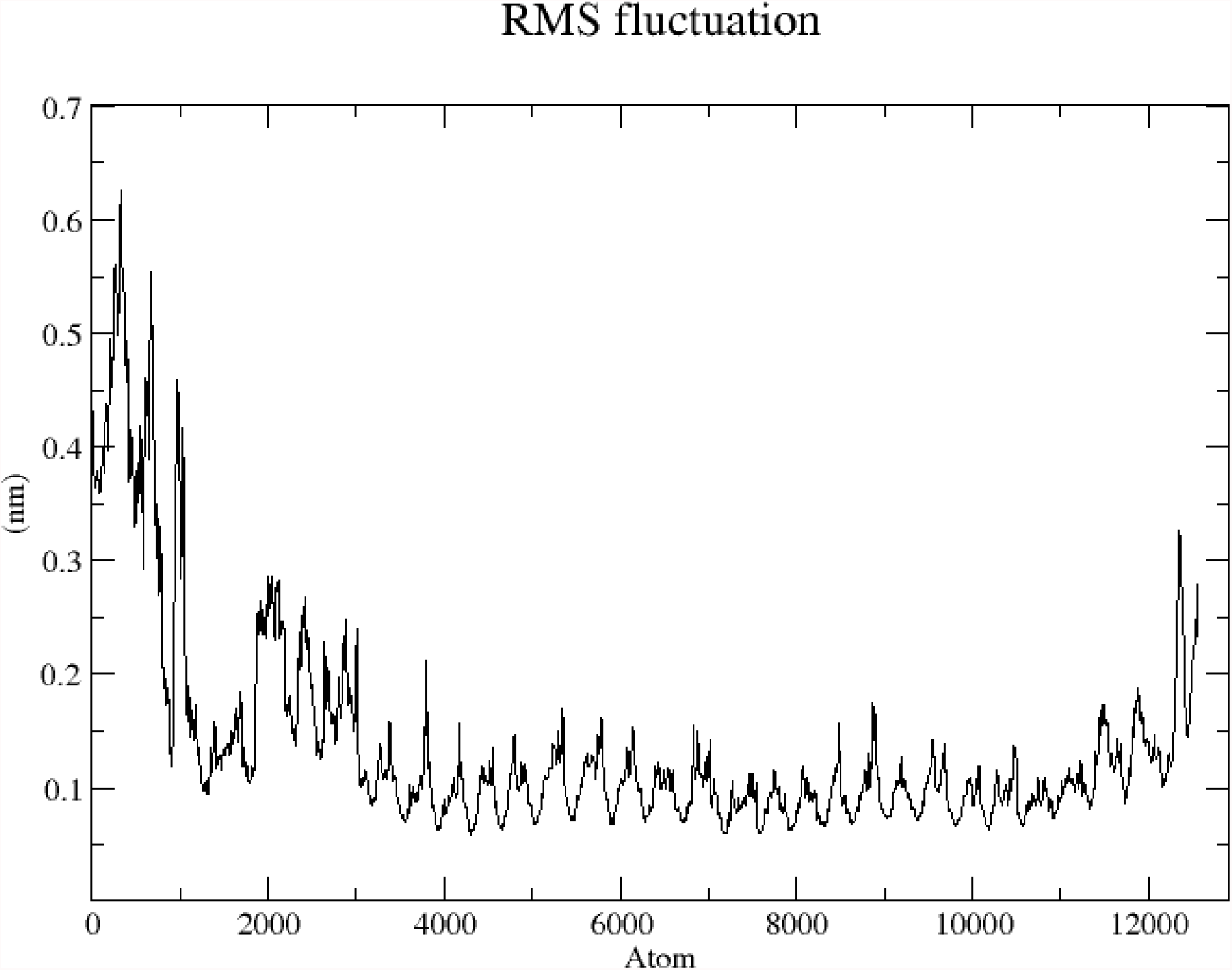
RMSF plot of the side chain residues of TLR 4-Vaccine Complex

### In Silico Cloning

The cloning and expression efficiency of the multi epitope vaccine construct in an expression vector was also determined by in-silico cloning. The Java Codon Adaptation Tool (JCat) was used for optimizing the codon usage of vaccine construct in *Escherichia coli* (strain K12). The optimized codon sequence was composed of 603 nucleotides. The GC content of the optimized nucleotide sequence of the multi epitope vaccine was 55.39%, which is well within the optimal GC percentage range (30% - 70%). The Codon Adaptation Index (CAI) was 0.964. The high CAI value indicates the possibility of efficient expression of the vaccine in the host (E. coli -strain K12). Finally, the SnapGene tool was utilized for insertion of the optimized codon sequences (multi epitope vaccine sequence) in pET28a (+) vector for restriction cloning design (Figure 10).

**Figure 10:**
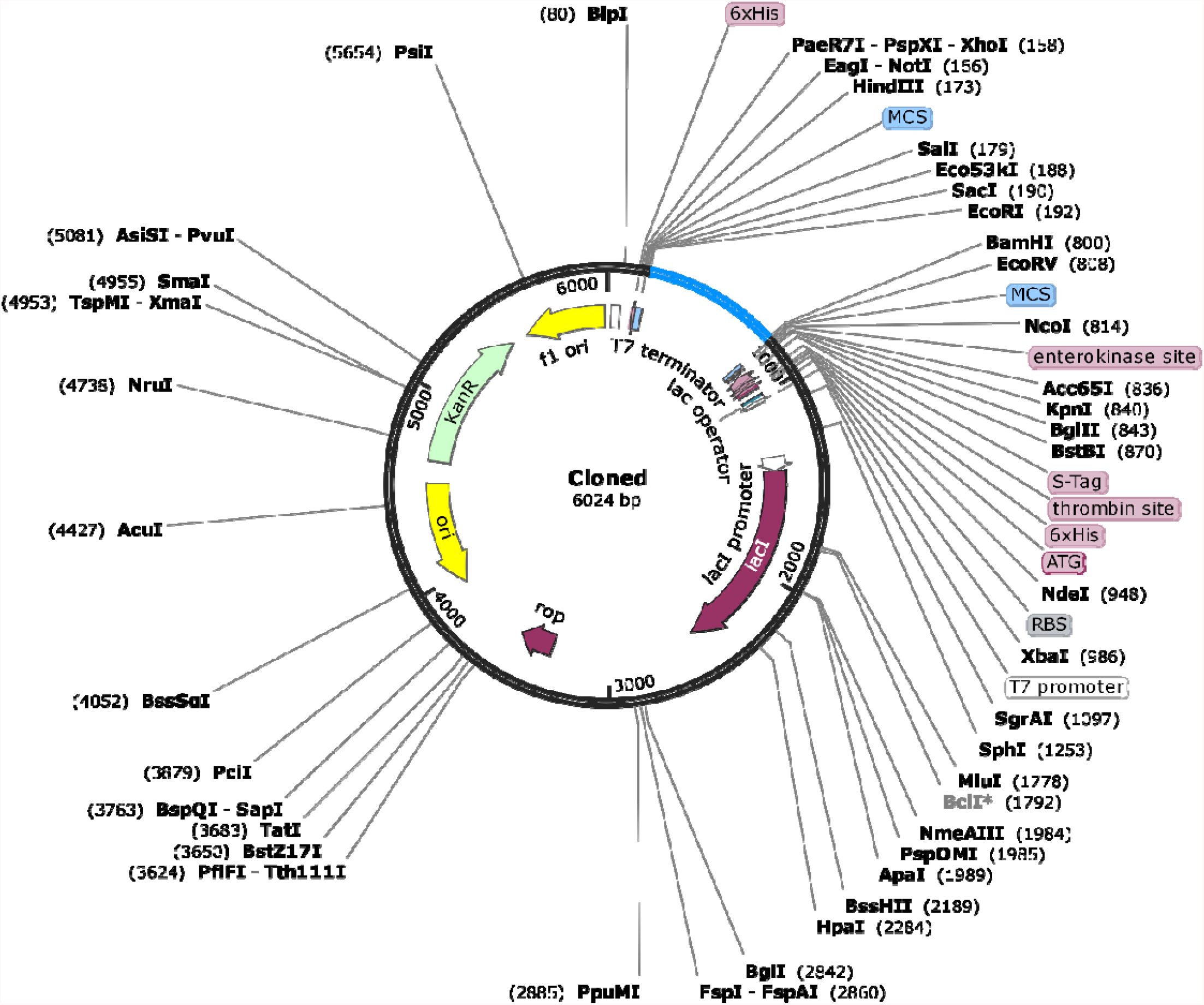
In silico restriction cloning design. The blue portion in the vector represents the multi epitope vaccine insert into the black pET28a(+) vector.

### Immune Simulation

C-ImmSim server immune simulation results show the prediction of the occurrence of primary immune response upon injection of the vaccine by showing high levels of IgM. The secondary and tertiary immune responses are also recorded by significant increase in B cell populations and in levels of IgG1 + IgG2, IgM, and IgG + IgM antibodies with a corresponding decrease in antigen concentration (Figure 11a). These results indicate the development of immune memory and consequently increased clearance of the antigen upon acceptance of consequent vaccine doses (Figure 11b). A similarly high response was seen in the T_H_ (helper) and T_C_ (cytotoxic) cell populations with corresponding memory development (Figure 11c, 11d). Repeated exposure with 3 injections (given at the intervals of four week) triggered increasing levels of IgG1 and decreasing levels of IgM while IFN- γ Concentration and T_H_ cell population were maintained at high levels throughout the duration of the vaccine injection.

**Figure 11a:**
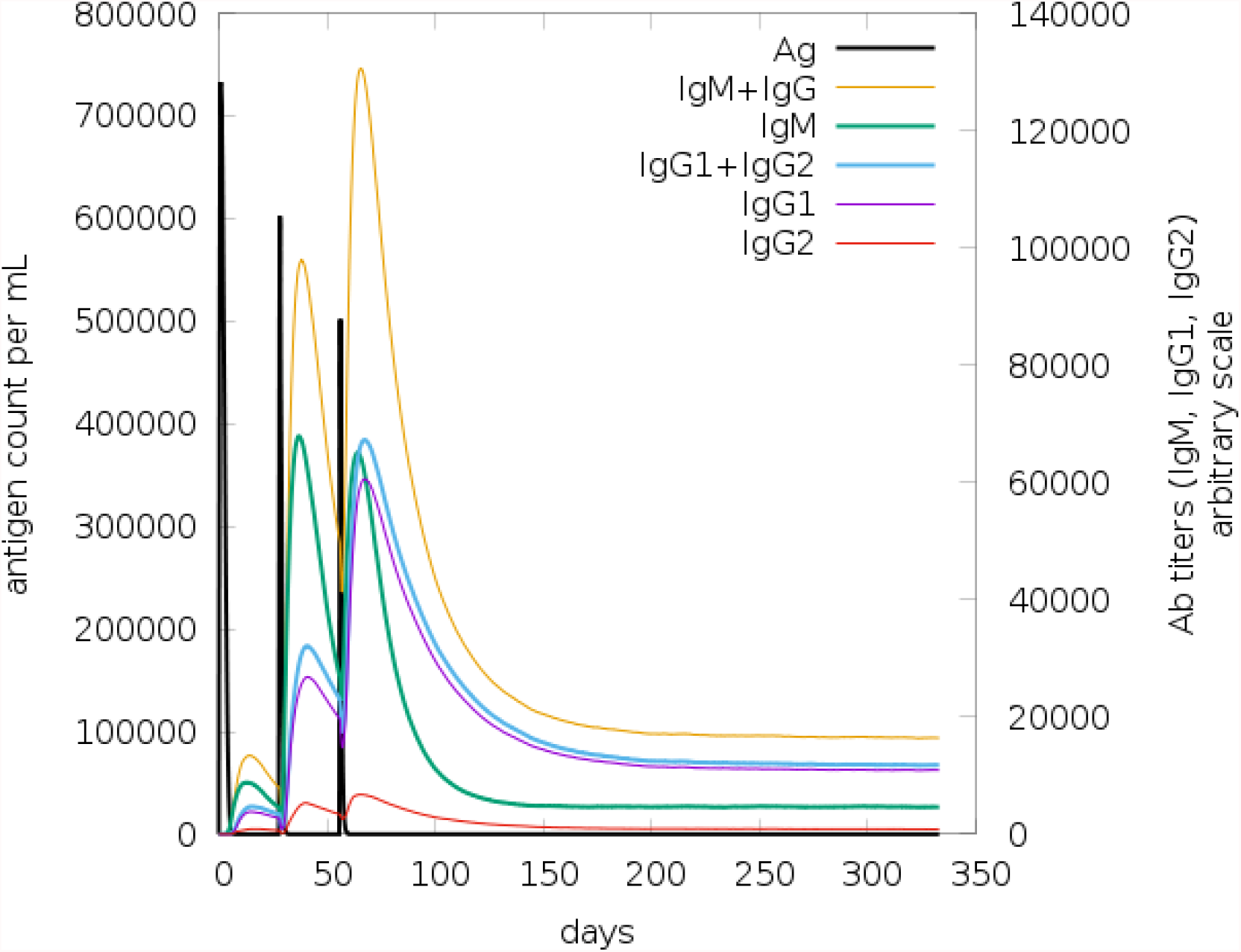
Immunoglobulin production in response to antigen injections (black vertical lines); specific subclasses are indicated as coloured peaks

**Figure 11b:**
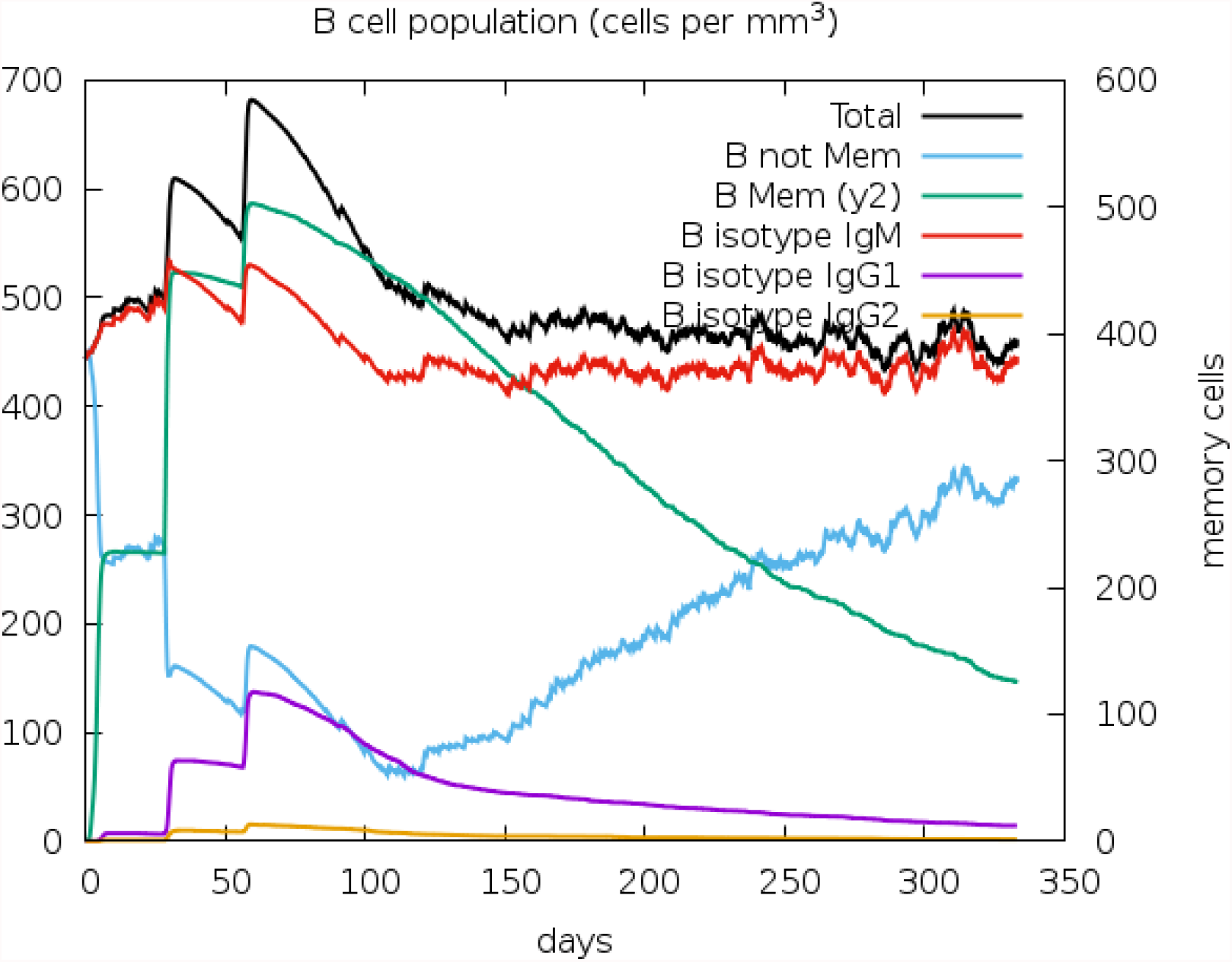
The evolution of B-cell populations after the three injections

**Figure 11c:**
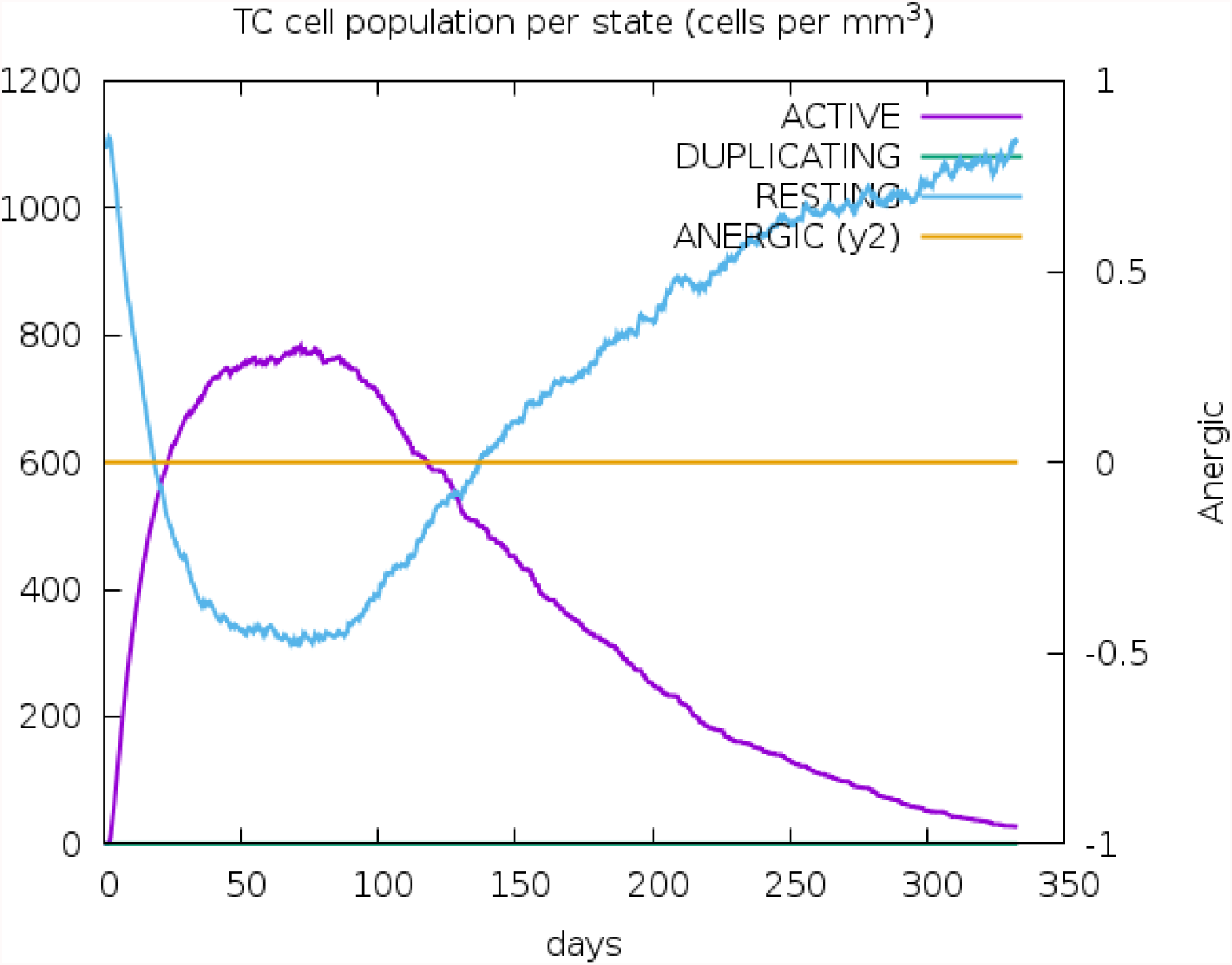
Cytotoxic T cell populations per state after the injections. The resting state represents cells not presented with the antigen while the anergic state represents tolerance of the T-cells to the antigen due to repeated exposures.

**Figure 11d:**
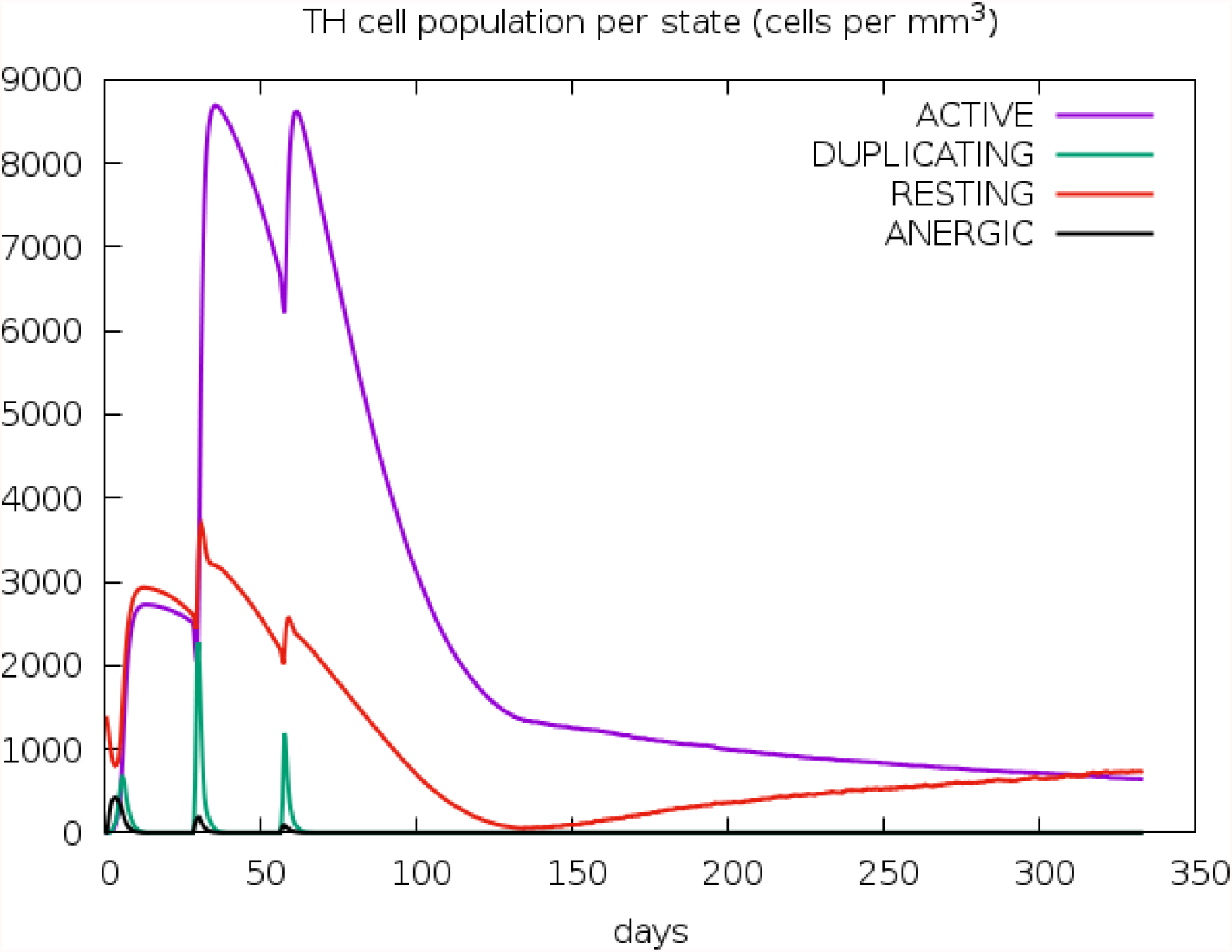
The evolution of Helper T cell populations per state after the injections.

## Discussion

Foodborne infection causes the death of a lot of people every year all over the world, especially in developing and underdeveloped countries (Newell et. al., 2010). *Salmonella* enterica serovar Typhi is the causative agent of typhoid disease, which is one of the most infamous foodborne illnesses. The goal of current study is to employ immunoinformatic techniques to build an efficient vaccine model that contains multiple *S*. Typhi epitopes. Multi epitope vaccines have some advantages over traditional vaccines due to having the following qualities: (a) it can be recognized by multiple T cell receptors; (b) it can activate both cellular and humoral immune responses, especially if it contains both T cell and B cell epitopes; (c) the vaccine’s multiple epitopes can be similar to epitopes of closely related pathogens, making it a broad spectrum vaccine; (d) a suitable adjuvant can be added to the vaccine to provide longevity as well as increased immunogenicity; and (e) by screening out the harmful epitopes, undesirable immune responses such autoimmunity and allergenicity can be prevented or greatly minimized.

The epitopes of the target proteins found in this study (T cell and IFN-epitopes) were chosen based on their ability to pass through multiple immune filters, including conservancy, promiscuity, localisation, antigenicity, allergenicity, and so on. Epitopes that failed to pass this screening method were excluded. Although Hemagglutinin was initially chosen for this investigation because of its better antigenicity score, all of the epitopes of it appeared to be too hazardous to be present in the multi epitope vaccine design.

If appropriate spacer sequences are placed between epitope areas, vaccine design can be improved (Meza et. al., 2017). AAY and GPGPG linkers were included among the anticipated epitopes in relevant researches (Ali et al., 2017, Khatoon et al., 2017), permitting the formulation of a powerful multi epitope immunization candidate (Meza et. al., 2017). The EAAAK linker was also placed between the adjuvant sequence and the peptide carrying the linked epitopes to guarantee a high level of expression and prolonged persistence of the vaccine molecules in the body, as previously described for bifunctional proteins (Arai et al., 2001).

Since *Salmonella enterica* is a gram-negative pathogen, the Toll like receptor-4 should be the key molecule to recognize it upon entry into the cell. Lipopolysaccharide (LPS), a major component of Gram-negative bacteria’s outer membrane, is effectively recognized by a cascade of LPS receptors and accessory proteins, including LPS binding protein (LBP), CD14, and the Toll-like receptor 4 (TLR-4)–MD-2 complex, as discussed in a study (Park et al., 2013). Another study (Hue et al. 2009) found that the amount of genetic diversity within TLR-4 in a Vietnamese community is linked to higher sensitivity to typhoidal infection, suggesting that TLR-4 may be involved in defence against typhoid fever.

Another critical stage in developing a multi epitope vaccine is effective cloning and expression in an appropriate expression vector. The value of the CAI (codon adaptation index) indicates codon usage bias. Although a CAI score of 1.0 is desirable, a score of >0.8 is regarded satisfactory (Morla et. al., 2016). A sequence’s GC content should be between 30 and 70 percent. Outside of this range, GC concentration has a negative impact on translational and transcriptional efficiency (Ali et. al., 2017). In order to ensure efficient expression and translation of the multi-epitope vaccine in an expression vector, pET-28a (+) expression vector was chosen. Several research groups have recently used a similar technique to build a multi-epitope vaccine against kaposi sarcoma, onchocerciasis, and other diseases (Chauhan et. al., 2019, Shey et. al., 2019). The multi epitope vaccine, which contains CTL, HTL, and IFN-epitopes, can activate several immune system components in humans, resulting in successful pathogen clearance. The effort to produce these epitopes contained at a single peptide in a bacterial system and execute the different immunological assays necessary to confirm the results acquired from in silico analysis would be the next step in this study’s research.

## Notes

### Competing Interest Statement

The authors have declared no competing interest.

