## Supplementary table 1-7 for "Designing a Multi-Epitope Vaccine Candidate against *Salmonella* Typhi Using Immunoinformatics Approach"

### **Supplementary Tables**

Supplementary Table 1: List of the whole genome sequence entries of Salmonella Typhi used in this study along with their strain names, IDs, size of genome, GC percentage and date of publication

| No. | Organism Name | Strain | Bio Sample ID | Size (Mb) | GC  % | Date Published |
| --- | --- | --- | --- | --- | --- | --- |
| 1 | Salmonella enterica subsp. enterica serovar Typhi  (S. Typhi) | CT18 | SAMEA1705914 | 5.13 | 51.88 | 07/11/2001 |
| 2 |  | Ty2 | SAMN02604095 | 4.79 | 52.10 | 21/03/2003 |
| 3 |  | P-stx-12 | SAMN02603101 | 4.95 | 51.89 | 01/02/2012 |
| 4 |  | Ty21a | SAMN02603210 | 4.79 | 52.00 | 01/05/2013 |
| 5 |  | STH2370 | SAMN02569978 | 4.76 | 52.10 | 27/01/2014 |
| 6 |  | PM016/13 | SAMN03765654 | 4.79 | 52.00 | 29/09/2015 |
| 7 |  | H12ESR04734-001A | SAMEA2072794 | 4.84 | 52.10 | 04/08/2017 |
| 8 |  | 343077_213147 | SAMN09320557 | 5.00 | 52.07 | 07/11/2018 |
| 9 |  | LXYSH | SAMN09630442 | 4.78 | 52.10 | 04/02/2019 |
| 10 |  | 4316STDY6559669 | SAMEA5577690 | 4.84 | 52.10 | 10/05/2019 |

Supplementary Table 2: List of potential T cell epitopes along with their position, amino acid sequence, interaction with HLA Class I and Class II alleles and supertypes, antigenicity and population coverage

| Protein Name | Position  (Amino acid no.) | Merged Peptide Sequence | HLA Class I alleles and supertypes | HLA Class II alleles | Antigenicity | Population coverage |
| --- | --- | --- | --- | --- | --- | --- |
| Peptidoglycan-associated lipoprotein | 127 | ANAVKMYLQGKGVSA | HLA-A*02, HLA-B*8, | HLA-DRB1*01:01 | 0.7754 | 51.79% |
|  | 71 | IVYFDLDKYDIRSDF | HLA-A*03,HLA-A*26,HLA-B*62 | HLA-DRB1*03:01 | 1.0921 | 36.01% |
|  | 80 | DIRSDFAAMLDAHAN | HLA-A*26,HLA-B*8,HLA-B*27,HLA-B*39 | HLA-DRB1*04:01 | 0.899 | 27.38% |
|  | 6 | VLKGLMIALPVMAIA | HLA-B*8,HLA-B*62 | HLA-DRB1*08:03 | 0.7188 | 12.60% |
|  | 62 | QMQQLQQNNIVYFDL | HLA-A*01,HLA-A*24,HLA-B*27,HLA-B*62,HLA-B*35:01 | HLA-DRB1*13:02 | 0.6069 | 29.37% |
| Outer Membrane Protein S_1_ | 5 | KVLALLVPALLVAGA | HLA-A*02,HLA-B*8,HLA-B*62,HLA-A*02:01 | HLA-DRB1*01:01 | 0.7079 | 51.79% |
|  | 207 | DGFGMSTSYDFDFGL | HLA-A*02,HLA-B*39,HLA-B*58,HLA-B*35:01,HLA-B*58:01 | HLA-DRB1*07:01 | 1.3532 | 57.29% |
|  | 308 | FGLRPSIAYLQSKGK | HLA-A*02,HLA-B*8,HLA-B*62,HLA-B*35:01 | HLA-DRB1*14:01 | 0.8875 | 52.81% |
|  | 143 | TDVYMLGRTNGVATY | HLA-A*02,HLA-B*8,HLA-B*62 | HLA-DRB1*04:01 | 0.4123 | 51.62% |
|  | 259 | KYDAYNVYLAAMYAE | HLA-A*02,HLA-A*24,HLA-B*39 | HLA-DRB1*15:01 | 0.6202 | 40.75% |
|  | 271 | VYLAAMYAETRNMTY | HLA-A*01,HLA-B*8,HLA-B*62,B35,HLA-B*35:01 | HLA-DRB1*04:01 | 0.4002 | 40.18% |
| Outer Membrane Protein F | 17 | PALLVAGAANAAEIY | HLA-A*01,HLA-B*58,HLA-B*62,HLA-B*35:01 | HLA-DRB1*01:01 | 0.5361 | 35.42% |
|  | 213 | DFDFGLSLGAAYSSS | HLA-A*26,HLA-B*8,HLA-B*62,HLA-B*35:01 | HLA-DRB1*01:01 | 1.094 | 32.09% |
|  | 205 | NGDGFGMSTSYDFDF | HLA-A*01,HLA-A*26,HLA-B*58,HLA-B*62,HLA-B*35:01,HLA-B*58:01 | HLA-DRB1*07:01 | 1.7599 | 31.66% |
|  | 258 | AKYDAYNVYLAAMYA | HLA-B*39,HLA-B*62 | HLA-DRB1*10:01,HLA-DRB1*15:01 | 0.6564 | 22.92% |
|  | 209 | YDFDFGLSLGAAYSS | HLA-B*39,HLA-B*44,HLA-B*35:01 | HLA-DRB1*01:01 | 1.168 | 21.30% |
|  | 359 | FNKNMSTYVDYKINL | HLA-A*24,HLA-B*39 | HLA-DRB1*07:01 | 0.7621 | 20.47% |
|  | 24 | AAEIYNKNGNKLDLY | HLA-A*24,HLA-B*39 | HLA-DRB1*13:02 | 1.4277 | 9.26% |
|  | 263 | AYNVYLAAMYAETRN | HLA-A*26,HLA-B*62 | HLA-DRB1*10:01 | 0.7825 | 8.26% |
|  | 94 | NSNLVRLAFAGLKYA | HLA-A*01,HLA-B*58,HLA-B*62,HLA-B*35:01 | HLA-DRB1*01:01,HLA-DRB1*12:01 | 0.5555 | 38.49% |
|  | 5 | KILAAVIPALLAAAT | HLA-A*B7,HLA-B*8,HLA-A*02:01 | HLA-DRB1*01:01,HLA-DRB1*08:03 | 0.4843 | 52.97% |
|  | 12 | PALLAAATANAAEIY | HLA-A*01,HLA-A*26,HLA-B*58,HLA-B*62 | HLA-DRB1*10:01,HLA-DRB1*08:03 | 0.5042 | 28.89% |
|  | 163 | DFFGLVDGLSFGIQY | HLA-B*58,HLA-B*62,HLA-B*35:01 | HLA-DRB1*07:01 | 0.8608 | 27.79% |
|  | 289 | LRPAISYVQSKGKQL | HLA-A*B7,HLA-B*8 | HLA-DRB1*07:01 | 0.4708 | 26.86% |
|  | 32 | KLDLYGKAVGRHVWT | HLA-B*39,HLA-B*58 | HLA-DRB1*07:01 | 0.8889 | 23.23% |
|  | 146 | DNYMTSRAGGLLTYR | HLA-A*26,HLA-B*27 | HLA-DRB1*07:01 | 0.4222 | 23.00% |
|  | 203 | FDGFGVTAAYSNSKR | HLA-A*26,HLA-B*62,HLA-B*35:01 | HLA-DRB1*10:01 | 0.6672 | 15.98% |
|  | 238 | AKYDANNVYLAAVYA | HLA-A*24,HLA-B*39,HLA-B*62 | HLA-DRB1*01:01 | 0.5344 | 13.96% |
|  | 283 | YQFDFGLRPAISYVQ | HLA-B*62,HLA-B*35:01 | HLA-DRB1*12:01 | 1.3319 | 12.51% |
| Hemagglutinin | 487 | GTGVKYIRTNDNGLE | HLA-A*26,HLA-B*8 | HLA-DRB1*04:01 | 1.4896 | 25.20% |
|  | 29 | KSGRRKLAVSALVGL | HLA-B*8,HLA-B*27,HLA-B*39 | HLA-DRB1*07:01 | 0.6109 | 28.99% |
|  | 360 | GSTSKITNVAAGALS | HLA-B*58,HLA-B*62 | HLA-DRB1*13:02 | 0.9189 | 9.89% |
|  | 181 | TKHHWEITNTFRYRI | HLA-A*26,HLA-B*44 | HLA-DRB1*07:01 | 1.2569 | 22.99% |
|  | 185 | WEITNTFRYRINEHW | HLA-B*27,HLA-B*58,HLA-B*62 | HLA-DRB1*07:01 | 1.073 | 21.04% |
| Outer Membrane Porin L | 199 | WLPYFELRWLDRNVG | HLA-B*58,HLA-B*62 | HLA-DRB1*11:01 | 1.262 | 13.60% |
|  | 150 | NYWNFIITDKFSYTF | HLA-A*01,HLA-A*24,HLA-A*26,HLA-B*39,HLA-B*58,HLA-B*62,HLA-B*58:01 | HLA-DRB1*12:01 | 1.1047 | 30.60% |
|  | 154 | FIITDKFSYTFEPHY | HLA-A*01,HLA-A*26,HLA-B*58,HLA-B*62,HLA-B*35:01,HLA-B*58:01 | HLA-DRB1*12:01 | 1.2504 | 34.72% |
|  | 216 | HREQNQIRIGAKYFF | HLA-A*26,HLA-B*27,HLA-B*39 | HLA-DRB1*12:01 | 1.0388 | 12.51% |
|  | 113 | KFTPWFNLTVRNRYN | HLA-A*01,HLA-A*02 | HLA-DRB1*14:01 | 0.9967 | 54.79% |
|  | 143 | NDSYEIGNYWNFIIT | HLA-B*58,HLA-B*62 | HLA-DRB1*15:01 | 0.569 | 21.20% |
|  | 146 | YEIGNYWNFIITDKF | HLA-A*24,HLA-A*26,HLA-B*44 | HLA-DRB1*15:01 | 0.4738 | 23.15% |

Supplementary Table 3: List of Potential IFN-Gamma Epitopes along with their potency score. The epitope sequences are listed with decreasing order of their score.

| Protein Name | Position  (Amino acid no.) | Epitope Sequence | Score (Threshold= score>0) |
| --- | --- | --- | --- |
| Peptidoglycan-associated lipoprotein | 143 | QISIVSYGKEKPAVL | 0.38556 |
|  | 144 | ISIVSYGKEKPAVLG | 0.30420 |
|  | 117 | EYNISLGERRANAVK | 0.30245 |
|  | 83 | SDFAAMLDAHANFLR | 0.27184 |
|  | 139 | VSADQISIVSYGKEK | 0.27039 |
|  | 85 | FAAMLDAHANFLRSN | 0.25025 |
|  | 82 | RSDFAAMLDAHANFL | 0.24894 |
|  | 90 | DAHANFLRSNPSYKV | 0.22907 |
|  | 149 | YGKEKPAVLGHDEAA | 0.22617 |
|  | 38 | LNGAGTGMDANGNGN | 0.20812 |
|  | 160 | DEAAYAKNRRAVLVY | 0.20018 |
|  | 88 | MLDAHANFLRSNPSY | 0.19880 |
|  | 142 | DQISIVSYGKEKPAV | 0.18813 |
|  | 118 | YNISLGERRANAVKM | 0.18570 |
|  | 135 | QGKGVSADQISIVSY | 0.17629 |
|  | 132 | MYLQGKGVSADQISI | 0.17054 |
|  | 137 | KGVSADQISIVSYGK | 0.16646 |
|  | 136 | GKGVSADQISIVSYG | 0.16646 |
|  | 86 | AAMLDAHANFLRSNP | 0.15410 |
|  | 133 | YLQGKGVSADQISIV | 0.13574 |
|  | 84 | DFAAMLDAHANFLRS | 0.13120 |
|  | 145 | SIVSYGKEKPAVLGH | 0.12051 |
|  | 89 | LDAHANFLRSNPSYK | 0.10919 |
|  | 107 | EGHADERGTPEYNIS | 0.10180 |
|  | 87 | AMLDAHANFLRSNPS | 0.10138 |
|  | 7 | LKGLMIALPVMAIAA | 0.09252 |
|  | 161 | EAAYAKNRRAVLVY | 0.09137 |
|  | 109 | HADERGTPEYNISLG | 0.08987 |
|  | 39 | NGAGTGMDANGNGNM | 0.08595 |
|  | 138 | GVSADQISIVSYGKE | 0.07850 |
|  | 111 | DERGTPEYNISLGER | 0.07691 |
|  | 91 | AHANFLRSNPSYKVT | 0.07581 |
|  | 113 | RGTPEYNISLGERRA | 0.07575 |
|  | 116 | PEYNISLGERRANAV | 0.06840 |
|  | 106 | VEGHADERGTPEYNI | 0.06227 |
|  | 166 | KNRRAVLVY | 0.05875 |
|  | 150 | GKEKPAVLGHDEAAY | 0.04849 |
|  | 148 | SYGKEKPAVLGHDEA | 0.04591 |
|  | 81 | IRSDFAAMLDAHANF | 0.01230 |
|  | 134 | LQGKGVSADQISIVS | 0.01203 |
|  | 8 | KGLMIALPVMAIAAC | 0.00026 |
| Outer Membrane Protein S_1_ | 309 | GLRPSIAYLQSKGKD | 1.09253 |
|  | 308 | FGLRPSIAYLQSKGK | 1.06308 |
|  | 386 | VGVGLVYQF | 0.96896 |
|  | 304 | YQFDFGLRPSIAYLQ | 0.94856 |
|  | 107 | YGSFDYGRNYGVIYD | 0.92724 |
|  | 307 | DFGLRPSIAYLQSKG | 0.87626 |
|  | 106 | EYGSFDYGRNYGVIY | 0.86894 |
|  | 303 | QYQFDFGLRPSIAYL | 0.85812 |
|  | 105 | GEYGSFDYGRNYGVI | 0.82852 |
|  | 302 | AQYQFDFGLRPSIAY | 0.82044 |
|  | 300 | VVAQYQFDFGLRPSI | 0.79642 |
|  | 301 | VAQYQFDFGLRPSIA | 0.78730 |
|  | 305 | QFDFGLRPSIAYLQS | 0.76211 |
|  | 2 | NRKVLALLVPALLVA | 0.74713 |
|  | 306 | FDFGLRPSIAYLQSK | 0.74536 |
|  | 278 | TYYGGGNGEGNGSIA | 0.72858 |
|  | 298 | FEVVAQYQFDFGLRP | 0.71677 |
|  | 179 | GGAGAGEGTGNGGNR | 0.70788 |
|  | 159 | FFGLVEGLNFALQYQ | 0.69776 |
|  | 3 | RKVLALLVPALLVAG | 0.69455 |
|  | 311 | RPSIAYLQSKGKDLG | 0.69192 |
|  | 279 | YYGGGNGEGNGSIAN | 0.69104 |
|  | 310 | LRPSIAYLQSKGKDL | 0.68600 |
|  | 108 | GSFDYGRNYGVIYDI | 0.67331 |
|  | 178 | NGGAGAGEGTGNGGN | 0.66800 |
|  | 4 | KVLALLVPALLVAGA | 0.65768 |
|  | 161 | GLVEGLNFALQYQGN | 0.65687 |
|  | 152 | ATYRNTDFFGLVEGL | 0.65628 |
|  | 299 | EVVAQYQFDFGLRPS | 0.65152 |
|  | 180 | GAGAGEGTGNGGNRK | 0.64258 |
|  | 277 | MTYYGGGNGEGNGSI | 0.64016 |
|  | 160 | FGLVEGLNFALQYQG | 0.63972 |
|  | 379 | GIATDDIVGVGLVYQ | 0.63194 |
|  | 5 | VLALLVPALLVAGAA | 0.62984 |
|  | 296 | QNFEVVAQYQFDFGL | 0.61440 |
|  | 111 | DYGRNYGVIYDIEAW | 0.60772 |
|  | 112 | YGRNYGVIYDIEAWT | 0.58438 |
|  | 77 | WEYNIKVNTTEGEGA | 0.56667 |
|  | 385 | IVGVGLVYQF | 0.55607 |
|  | 164 | EGLNFALQYQGNNEN | 0.55512 |
|  | 104 | FGEYGSFDYGRNYGV | 0.55304 |
|  | 157 | TDFFGLVEGLNFALQ | 0.54820 |
|  | 297 | NFEVVAQYQFDFGLR | 0.53620 |
|  | 158 | DFFGLVEGLNFALQY | 0.50691 |
|  | 110 | FDYGRNYGVIYDIEA | 0.50573 |
|  | 76 | QWEYNIKVNTTEGEG | 0.49924 |
|  | 280 | YGGGNGEGNGSIANK | 0.49776 |
|  | 380 | IATDDIVGVGLVYQF | 0.49275 |
|  | 275 | RNMTYYGGGNGEGNG | 0.49137 |
|  | 109 | SFDYGRNYGVIYDIE | 0.49081 |
|  | 156 | NTDFFGLVEGLNFAL | 0.48829 |
|  | 6 | LALLVPALLVAGAAN | 0.48707 |
|  | 381 | ATDDIVGVGLVYQF | 0.47652 |
|  | 155 | RNTDFFGLVEGLNFA | 0.46469 |
|  | 165 | GLNFALQYQGNNENG | 0.46006 |
|  | 78 | EYNIKVNTTEGEGAN | 0.44883 |
|  | 177 | ENGGAGAGEGTGNGG | 0.44844 |
|  | 384 | DIVGVGLVYQF | 0.44672 |
|  | 113 | GRNYGVIYDIEAWTD | 0.44574 |
|  | 283 | GNGEGNGSIANKTQN | 0.43115 |
|  | 154 | YRNTDFFGLVEGLNF | 0.42982 |
|  | 1 | MNRKVLALLVPALLV | 0.42898 |
|  | 153 | TYRNTDFFGLVEGLN | 0.41463 |
|  | 172 | YQGNNENGGAGAGEG | 0.40329 |
|  | 274 | TRNMTYYGGGNGEGN | 0.40177 |
|  | 382 | TDDIVGVGLVYQF | 0.39088 |
|  | 115 | NYGVIYDIEAWTDAL | 0.35948 |
|  | 276 | NMTYYGGGNGEGNGS | 0.35303 |
|  | 103 | KFGEYGSFDYGRNYG | 0.34913 |
|  | 312 | PSIAYLQSKGKDLGG | 0.33143 |
|  | 81 | IKVNTTEGEGANSWT | 0.32978 |
|  | 163 | VEGLNFALQYQGNNE | 0.31525 |
|  | 383 | DDIVGVGLVYQF | 0.30217 |
|  | 294 | KTQNFEVVAQYQFDF | 0.30093 |
|  | 324 | LGGQEVHRGNWRYTD | 0.29999 |
|  | 323 | DLGGQEVHRGNWRYT | 0.29922 |
|  | 114 | RNYGVIYDIEAWTDA | 0.29301 |
|  | 281 | GGGNGEGNGSIANKT | 0.28578 |
|  | 181 | AGAGEGTGNGGNRKL | 0.28027 |
|  | 151 | VATYRNTDFFGLVEG | 0.27731 |
|  | 284 | NGEGNGSIANKTQNF | 0.27268 |
|  | 79 | YNIKVNTTEGEGANS | 0.26614 |
|  | 80 | NIKVNTTEGEGANSW | 0.26036 |
|  | 289 | GSIANKTQNFEVVAQ | 0.25683 |
|  | 378 | NGIATDDIVGVGLVY | 0.25636 |
|  | 295 | TQNFEVVAQYQFDFG | 0.24772 |
|  | 286 | EGNGSIANKTQNFEV | 0.23843 |
|  | 377 | NNGIATDDIVGVGLV | 0.22020 |
|  | 282 | GGNGEGNGSIANKTQ | 0.20981 |
|  | 116 | YGVIYDIEAWTDALP | 0.18433 |
|  | 171 | QYQGNNENGGAGAGE | 0.18414 |
|  | 174 | GNNENGGAGAGEGTG | 0.17665 |
|  | 175 | NNENGGAGAGEGTGN | 0.17665 |
|  | 206 | STSYDFDFGLSLGAA | 0.16887 |
|  | 285 | GEGNGSIANKTQNFE | 0.16863 |
|  | 162 | LVEGLNFALQYQGNN | 0.16672 |
|  | 287 | GNGSIANKTQNFEVV | 0.15990 |
|  | 7 | ALLVPALLVAGAANA | 0.15353 |
|  | 74 | YGQWEYNIKVNTTEG | 0.14176 |
|  | 371 | DDDFYANNGIATDDI | 0.14095 |
|  | 170 | LQYQGNNENGGAGAG | 0.14017 |
|  | 321 | GKDLGGQEVHRGNWR | 0.13624 |
|  | 176 | NENGGAGAGEGTGNG | 0.13612 |
|  | 293 | NKTQNFEVVAQYQFD | 0.13272 |
|  | 322 | KDLGGQEVHRGNWRY | 0.13262 |
|  | 8 | LLVPALLVAGAANAA | 0.13119 |
|  | 214 | GLSLGAAYSSSDRSD | 0.12645 |
|  | 75 | GQWEYNIKVNTTEGE | 0.12367 |
|  | 319 | SKGKDLGGQEVHRGN | 0.11797 |
|  | 188 | GNGGNRKLARENGDG | 0.10943 |
|  | 101 | GLKFGEYGSFDYGRN | 0.10916 |
|  | 167 | NFALQYQGNNENGGA | 0.10646 |
|  | 325 | GGQEVHRGNWRYTDK | 0.10426 |
|  | 17 | GAANAAEIYNKNGNK | 0.10048 |
|  | 150 | GVATYRNTDFFGLVE | 0.09256 |
|  | 290 | SIANKTQNFEVVAQY | 0.09024 |
|  | 291 | IANKTQNFEVVAQYQ | 0.08857 |
|  | 187 | TGNGGNRKLARENGD | 0.08531 |
|  | 205 | MSTSYDFDFGLSLGA | 0.08498 |
|  | 328 | EVHRGNWRYTDKDLV | 0.07562 |
|  | 169 | ALQYQGNNENGGAGA | 0.07130 |
|  | 313 | SIAYLQSKGKDLGGQ | 0.06903 |
|  | 117 | GVIYDIEAWTDALPE | 0.06564 |
|  | 376 | ANNGIATDDIVGVGL | 0.06545 |
|  | 173 | QGNNENGGAGAGEGT | 0.06286 |
|  | 182 | GAGEGTGNGGNRKLA | 0.05719 |
|  | 292 | ANKTQNFEVVAQYQF | 0.05002 |
|  | 185 | EGTGNGGNRKLAREN | 0.04005 |
|  | 207 | TSYDFDFGLSLGAAY | 0.03992 |
|  | 370 | EDDDFYANNGIATDD | 0.03971 |
|  | 102 | LKFGEYGSFDYGRNY | 0.03652 |
|  | 330 | HRGNWRYTDKDLVKY | 0.03605 |
|  | 166 | LNFALQYQGNNENGG | 0.03444 |
|  | 320 | KGKDLGGQEVHRGNW | 0.02581 |
|  | 204 | GMSTSYDFDFGLSLG | 0.01801 |
|  | 220 | AYSSSDRSDNQVARG | 0.01740 |
|  | 254 | WTIGAKYDAYNVYLA | 0.01408 |
| Outer Membrane Protein F | 6 | ILAAVIPALLAAATA | 0.97912 |
|  | 278 | EVVAQYQFDFGLRPA | 0.78227 |
|  | 277 | LEVVAQYQFDFGLRP | 0.77648 |
|  | 5 | KILAAVIPALLAAAT | 0.77303 |
|  | 7 | LAAVIPALLAAATAN | 0.74206 |
|  | 4 | RKILAAVIPALLAAA | 0.72052 |
|  | 280 | VAQYQFDFGLRPAIS | 0.67466 |
|  | 153 | AGGLLTYRNSDFFGL | 0.65317 |
|  | 348 | YVGTDDQAAVGITYQ | 0.61919 |
|  | 151 | SRAGGLLTYRNSDFF | 0.58390 |
|  | 283 | YQFDFGLRPAISYVQ | 0.58031 |
|  | 287 | FGLRPAISYVQSKGK | 0.57962 |
|  | 279 | VVAQYQFDFGLRPAI | 0.57689 |
|  | 355 | AAVGITYQF | 0.57374 |
|  | 346 | SSYVGTDDQAAVGIT | 0.53582 |
|  | 345 | SSSYVGTDDQAAVGI | 0.53247 |
|  | 8 | AAVIPALLAAATANA | 0.49255 |
|  | 9 | AVIPALLAAATANAA | 0.49255 |
|  | 154 | GGLLTYRNSDFFGLV | 0.49230 |
|  | 3 | KRKILAAVIPALLAA | 0.47707 |
|  | 349 | VGTDDQAAVGITYQF | 0.47678 |
|  | 281 | AQYQFDFGLRPAISY | 0.47493 |
|  | 282 | QYQFDFGLRPAISYV | 0.46507 |
|  | 152 | RAGGLLTYRNSDFFG | 0.46405 |
|  | 288 | GLRPAISYVQSKGKQ | 0.46278 |
|  | 284 | QFDFGLRPAISYVQS | 0.46216 |
|  | 275 | QNLEVVAQYQFDFGL | 0.42940 |
|  | 347 | SYVGTDDQAAVGITY | 0.42547 |
|  | 286 | DFGLRPAISYVQSKG | 0.42328 |
|  | 78 | QWEYRTKADRAEGEQ | 0.42082 |
|  | 229 | DRAESWAVGAKYDAN | 0.40937 |
|  | 2 | MKRKILAAVIPALLA | 0.40111 |
|  | 150 | TSRAGGLLTYRNSDF | 0.39510 |
|  | 80 | EYRTKADRAEGEQQN | 0.38676 |
|  | 285 | FDFGLRPAISYVQSK | 0.37446 |
|  | 79 | WEYRTKADRAEGEQQ | 0.37337 |
|  | 155 | GLLTYRNSDFFGLVD | 0.36770 |
|  | 276 | NLEVVAQYQFDFGLR | 0.36660 |
|  | 77 | GQWEYRTKADRAEGE | 0.35861 |
|  | 230 | RAESWAVGAKYDANN | 0.34080 |
|  | 142 | GAYTDNYMTSRAGGL | 0.32110 |
|  | 34 | DLYGKAVGRHVWTTT | 0.31782 |
|  | 146 | DNYMTSRAGGLLTYR | 0.29254 |
|  | 85 | ADRAEGEQQNSNLVR | 0.28236 |
|  | 82 | RTKADRAEGEQQNSN | 0.27941 |
|  | 354 | QAAVGITYQF | 0.25991 |
|  | 86 | DRAEGEQQNSNLVRL | 0.25887 |
|  | 38 | KAVGRHVWTTTGDSK | 0.24804 |
|  | 157 | LTYRNSDFFGLVDGL | 0.23531 |
|  | 15 | LAAATANAAEIYNKD | 0.22482 |
|  | 1 | MMKRKILAAVIPALL | 0.22339 |
|  | 76 | FGQWEYRTKADRAEG | 0.21147 |
|  | 143 | AYTDNYMTSRAGGLL | 0.21101 |
|  | 14 | LLAAATANAAEIYNK | 0.20236 |
|  | 81 | YRTKADRAEGEQQNS | 0.20052 |
|  | 228 | GDRAESWAVGAKYDA | 0.19666 |
|  | 145 | TDNYMTSRAGGLLTY | 0.19660 |
|  | 289 | LRPAISYVQSKGKQL | 0.18787 |
|  | 309 | SADLAKYIQAGATYY | 0.18632 |
|  | 147 | NYMTSRAGGLLTYRN | 0.17670 |
|  | 35 | LYGKAVGRHVWTTTG | 0.17393 |
|  | 84 | KADRAEGEQQNSNLV | 0.17224 |
|  | 16 | AAATANAAEIYNKDG | 0.16614 |
|  | 226 | GNGDRAESWAVGAKY | 0.16449 |
|  | 149 | MTSRAGGLLTYRNSD | 0.16342 |
|  | 10 | VIPALLAAATANAAE | 0.15912 |
|  | 39 | AVGRHVWTTTGDSKN | 0.15451 |
|  | 344 | YSSSYVGTDDQAAVG | 0.14553 |
|  | 148 | YMTSRAGGLLTYRNS | 0.14501 |
|  | 273 | KTQNLEVVAQYQFDF | 0.14431 |
|  | 106 | KYAEVGSIDYGRNYG | 0.13490 |
|  | 37 | GKAVGRHVWTTTGDS | 0.13359 |
|  | 177 | YQGKNQDNHSINSQN | 0.11543 |
|  | 141 | GGAYTDNYMTSRAGG | 0.11503 |
|  | 103 | AGLKYAEVGSIDYGR | 0.11465 |
|  | 233 | SWAVGAKYDANNVYL | 0.10979 |
|  | 231 | AESWAVGAKYDANNV | 0.10337 |
|  | 102 | FAGLKYAEVGSIDYG | 0.09997 |
|  | 144 | YTDNYMTSRAGGLLT | 0.09973 |
|  | 290 | RPAISYVQSKGKQLN | 0.09737 |
|  | 165 | FGLVDGLSFGIQYQG | 0.09704 |
|  | 227 | NGDRAESWAVGAKYD | 0.09181 |
|  | 36 | YGKAVGRHVWTTTGD | 0.08743 |
|  | 13 | ALLAAATANAAEIYN | 0.08201 |
|  | 274 | TQNLEVVAQYQFDFG | 0.07957 |
|  | 232 | ESWAVGAKYDANNVY | 0.06789 |
|  | 234 | WAVGAKYDANNVYLA | 0.06685 |
|  | 83 | TKADRAEGEQQNSNL | 0.06621 |
|  | 305 | GADGSADLAKYIQAG | 0.06434 |
|  | 306 | ADGSADLAKYIQAGA | 0.06434 |
|  | 353 | DQAAVGITYQF | 0.06298 |
|  | 104 | GLKYAEVGSIDYGRN | 0.05939 |
|  | 164 | FFGLVDGLSFGIQYQ | 0.05651 |
|  | 225 | DGNGDRAESWAVGAK | 0.05639 |
|  | 107 | YAEVGSIDYGRNYGI | 0.04804 |
|  | 270 | MANKTQNLEVVAQYQ | 0.03446 |
|  | 308 | GSADLAKYIQAGATY | 0.02499 |
|  | 350 | GTDDQAAVGITYQF | 0.02197 |
|  | 310 | ADLAKYIQAGATYYF | 0.01858 |
|  | 75 | GFGQWEYRTKADRAE | 0.01164 |
|  | 351 | TDDQAAVGITYQF | 0.00436 |
| Outer Membrane Porin L | 6 | TLVILTSVISTSVFA | 0.63007 |
|  | 5 | NTLVILTSVISTSVF | 0.47403 |
|  | 7 | LVILTSVISTSVFAG | 0.44929 |
|  | 1 | MKSLNTLVILTSVIS | 0.42489 |
|  | 2 | KSLNTLVILTSVIST | 0.42258 |
|  | 91 | PGGLINDKSIGSGGA | 0.35976 |
|  | 12 | SVISTSVFAGAYVEN | 0.35301 |
|  | 92 | GGLINDKSIGSGGAV | 0.35268 |
|  | 11 | TSVISTSVFAGAYVE | 0.32712 |
|  | 4 | LNTLVILTSVISTSV | 0.31617 |
|  | 10 | LTSVISTSVFAGAYV | 0.27972 |
|  | 13 | VISTSVFAGAYVENR | 0.27553 |
|  | 66 | ELKHGYNEIEGWYPL | 0.27112 |
|  | 203 | FELRWLDRNVGLYHR | 0.25504 |
|  | 181 | TKHHWEITNTFRYRI | 0.22861 |
|  | 3 | SLNTLVILTSVISTS | 0.22536 |
|  | 8 | VILTSVISTSVFAGA | 0.20445 |
|  | 68 | KHGYNEIEGWYPLFK | 0.19760 |
|  | 93 | GLINDKSIGSGGAVY | 0.18594 |
|  | 90 | QPGGLINDKSIGSGG | 0.16547 |
|  | 84 | TDKLTIQPGGLINDK | 0.15540 |
|  | 34 | SDQMEFMLRVGYNSD | 0.13713 |
|  | 9 | ILTSVISTSVFAGAY | 0.13096 |
|  | 69 | HGYNEIEGWYPLFKP | 0.12907 |
|  | 202 | YFELRWLDRNVGLYH | 0.11073 |
|  | 65 | DELKHGYNEIEGWYP | 0.10300 |
|  | 64 | DDELKHGYNEIEGWY | 0.10061 |
|  | 17 | SVFAGAYVENREAYN | 0.09922 |
|  | 204 | ELRWLDRNVGLYHRE | 0.09808 |
|  | 85 | DKLTIQPGGLINDKS | 0.08516 |
|  | 67 | LKHGYNEIEGWYPLF | 0.08363 |
|  | 184 | HWEITNTFRYRINEH | 0.06282 |
|  | 185 | WEITNTFRYRINEHW | 0.06282 |
|  | 182 | KHHWEITNTFRYRIN | 0.06072 |
|  | 32 | LASDQMEFMLRVGYN | 0.05681 |
|  | 194 | RINEHWLPYFELRWL | 0.04709 |
|  | 197 | EHWLPYFELRWLDRN | 0.04374 |
|  | 70 | GYNEIEGWYPLFKPT | 0.04024 |
|  | 198 | HWLPYFELRWLDRNV | 0.03959 |
|  | 180 | GTKHHWEITNTFRYR | 0.03605 |
|  | 33 | ASDQMEFMLRVGYNS | 0.02289 |
|  | 16 | TSVFAGAYVENREAY | 0.01670 |
|  | 205 | LRWLDRNVGLYHREQ | 0.00509 |
|  | 183 | HHWEITNTFRYRINE | 0.00282 |
|  | 73 | EIEGWYPLFKPTDKL | 0.00087 |
|  | 72 | NEIEGWYPLFKPTDK | 0.00075 |

Supplementary Table 4: Similarity Search of Selected Epitopes among Other *Salmonella* Pathogenic Serovars:

| Protein Name | Merged Peptide Sequence | Serovar | Query Cover | E Value | Percent Identity |
| --- | --- | --- | --- | --- | --- |
| Peptidoglycan-associated lipoprotein | ANAVKMYLQGKGVSA | *Salmonella* Paratyphi A (taxid:54388)  *Salmonella* Typhimurium (taxid:90371)  *Salmonella* Enteritidis (taxid:149539) | 100%  100%  100% | 2e-09  4e-08  3e-08 | 100.00%  100.00%  100.00% |
| Outer Membrane Protein S_1_ | KVLALLVPALLVAGA | *Salmonella* Paratyphi A (taxid:54388)  *Salmonella* Typhimurium (taxid:90371)  *Salmonella* Enteritidis (taxid:149539) | 100%  100%  100% | 3e-08  7e-08  9e-08 | 100.00%  100.00%  100.00% |
|  | DGFGMSTSYDFDFGL | *Salmonella* Paratyphi A (taxid:54388)  *Salmonella* Typhimurium (taxid:90371)  *Salmonella* Enteritidis (taxid:149539) | 100%  100%  100% | 2e-10  3e-09  2e-09 | 100.00%  100.00%  100.00% |
|  | FGLRPSIAYLQSKGK | *Salmonella* Paratyphi A (taxid:54388)  *Salmonella* Typhimurium (taxid:90371)  *Salmonella* Enteritidis (taxid:149539) | 100%  100%  100% | 2e-09  3e-08  2e-08 | 100.00%  100.00%  100.00% |

| Protein Name | Merged Peptide Sequence | Serovar | Query Cover | E Value | Percent Identity |
| --- | --- | --- | --- | --- | --- |
| Outer Membrane Protein S_1_ | TDVYMLGRTNGVATY | *Salmonella* Paratyphi A (taxid:54388)  *Salmonella* Typhimurium (taxid:90371)  *Salmonella* Enteritidis (taxid:149539) | 100%  100%  100% | 2e-10  4e-09  3e-09 | 100.00%  100.00%  100.00% |
|  | FGLRPSIAYLQSKGK | *Salmonella* Paratyphi A (taxid:54388)  *Salmonella* Typhimurium (taxid:90371)  *Salmonella* Enteritidis (taxid:149539) | 100%  100%  100% | 2e-09  3e-08  2e-08 | 100.00%  100.00%  100.00% |
|  | QFDFGLRPSIAYLQS | *Salmonella* Paratyphi A (taxid:54388)  *Salmonella* Typhimurium (taxid:90371)  *Salmonella* Enteritidis (taxid:149539) | 100%  100%  100% | 3e-10  5e-09  4e-09 | 100.00%  100.00%  100.00% |
|  | YQFDFGLRPSIAYLQ | *Salmonella* Paratyphi A (taxid:54388)  *Salmonella* Typhimurium (taxid:90371)  *Salmonella* Enteritidis (taxid:149539) | 100%  100%  100% | 8e-11  1e-09  1e-09 | 100.00%  100.00%  100.00% |
| Outer Membrane Protein F | KILAAVIPALLAAAT | *Salmonella* Paratyphi A (taxid:54388)  *Salmonella* Typhimurium (taxid:90371)  *Salmonella* Enteritidis (taxid:149539) | 100%  100%  100% | 2e-08  3e-07  2e-07 | 100.00%  100.00%  100.00% |
|  | ILAAVIPALLAAATA | *Salmonella* Paratyphi A (taxid:54388)  *Salmonella* Typhimurium (taxid:90371)  *Salmonella* Enteritidis (taxid:149539) | 100%  100%  100% | 3e-08  2e-07  2e-07 | 100.00%  100.00%  100.00% |
| Outer Membrane Porin L | KFTPWFNLTVRNRYN | *Salmonella* Paratyphi A (taxid:54388)  *Salmonella* Typhimurium (taxid:90371)  *Salmonella* Enteritidis (taxid:149539) | 100%  100%  100% | 1e-11  2e-10  1e-10 | 100.00%  100.00%  100.00% |

Supplementary Table 5: Structure Information of Refined Models by Galaxy Refine Server 2.

| Model | RMSD | MolProbity | Clash score | Poor rotamers | Rama favored | GALAXY energy |
| --- | --- | --- | --- | --- | --- | --- |
| Initial | 0.000 | 3.728 | 49.1 | 12.9 | 81.4 | 540.46 |
| MODEL 1 | 3.816 | 1.201 | 0.9 | 0.0 | 93.5 | -3494.15 |
| MODEL 2 | 2.747 | 1.261 | 0.9 | 0.7 | 92.0 | -3474.86 |
| MODEL 3 | 3.825 | 1.340 | 1.2 | 0.0 | 91.5 | -3471.87 |
| MODEL 4 | 2.789 | 1.222 | 0.9 | 0.7 | 93.0 | -3466.26 |
| MODEL 5 | 3.229 | 1.201 | 0.9 | 0.7 | 93.5 | -3465.54 |
| MODEL 6 | 2.697 | 1.112 | 1.2 | 0.0 | 96.0 | -3459.76 |
| MODEL 7 | 2.951 | 1.323 | 1.2 | 0.0 | 92.0 | -3459.10 |
| MODEL 8 | 3.065 | 1.213 | 1.2 | 0.0 | 94.5 | -3458.43 |
| MODEL 9 | 3.167 | 1.323 | 1.2 | 0.0 | 92.0 | -3452.03 |
| MODEL 10 | 1.661 | 1.116 | 0.3 | 0.0 | 91.5 | -3441.79 |

Supplementary table 6: List of protein-protein docking pose scores

| Title | PIPER pose energy | PIPER pose score | PIPER Model Number | PIPER Cluster Size | Potential Energy-OPLS3e |
| --- | --- | --- | --- | --- | --- |
| prot-prot-docking_2_pose_1 | -1251.42 | -197.798 | 6 | 19 | 16351.15 |
| prot-prot-docking_2_pose_2 | -1238.5 | -209.297 | 8 | 17 | 12218.52 |
| prot-prot-docking_2_pose_3 | -1222.64 | -12.347 | 10 | 13 | 15078.89 |
| prot-prot-docking_2_pose_4 | -1165.06 | -330.118 | 30 | 14 | 17568.55 |
| prot-prot-docking_2_pose_5 | -1154.53 | -20.33 | 36 | 26 | 13201.24 |
| prot-prot-docking_2_pose_6 | -1152.57 | -277.814 | 37 | 16 | 17911.62 |
| prot-prot-docking_2_pose_7 | -1151.27 | -77.692 | 38 | 17 | 13650.9 |
| prot-prot-docking_2_pose_8 | -1145.2 | -181.587 | 44 | 20 | 14293.26 |
| prot-prot-docking_2_pose_9 | -1134.91 | -428.929 | 58 | 18 | 16108.93 |
| prot-prot-docking_2_pose_10 | -1125.71 | -384.724 | 66 | 15 | 16645.12 |
| prot-prot-docking_2_pose_11 | -1119.95 | -471.603 | 79 | 20 | 21641.06 |
| prot-prot-docking_2_pose_12 | -1107.92 | -339.058 | 106 | 14 | 18276.49 |
| prot-prot-docking_2_pose_13 | -1097.66 | -77.129 | 123 | 13 | 15648.81 |
| prot-prot-docking_2_pose_14 | -1088.59 | 85.586 | 140 | 12 | 11563.6 |
| prot-prot-docking_2_pose_15 | -1087.35 | -165.858 | 144 | 19 | 18846.96 |
| prot-prot-docking_2_pose_16 | -1083.85 | -347.682 | 157 | 17 | 21946.74 |
| prot-prot-docking_2_pose_17 | -1081.64 | -248.469 | 167 | 18 | 13370.42 |
| prot-prot-docking_2_pose_18 | -1056.5 | -203.836 | 240 | 11 | 16972.78 |
| prot-prot-docking_2_pose_19 | -1049.68 | -567.63 | 270 | 19 | 12418.19 |
| prot-prot-docking_2_pose_20 | -1039.54 | -432.571 | 326 | 16 | 20340.76 |
| prot-prot-docking_2_pose_21 | -1023.11 | 18.561 | 420 | 15 | 17452.88 |
| prot-prot-docking_2_pose_22 | -1014.29 | -326.437 | 493 | 9 | 22686.94 |
| prot-prot-docking_2_pose_23 | -1007.84 | -360.683 | 549 | 23 | 16228.13 |
| prot-prot-docking_2_pose_24 | -1006.85 | -120.164 | 562 | 15 | 19101.06 |
| prot-prot-docking_2_pose_25 | -996.804 | -113.879 | 673 | 34 | 14029.69 |
| prot-prot-docking_2_pose_26 | -982.213 | -246.714 | 850 | 7 | 18792.81 |
| prot-prot-docking_2_pose_27 | -980.321 | -42.988 | 876 | 8 | 18183.96 |
| prot-prot-docking_2_pose_28 | -979.466 | -29.472 | 886 | 19 | 13458.16 |
| prot-prot-docking_2_pose_29 | -978.803 | -251.97 | 898 | 23 | 17054.98 |
| prot-prot-docking_2_pose_30 | -973.697 | -199.677 | 978 | 23 | 21845.3 |

Supplementary Table 7: Interactions between the residues of TLR-4 protein and the designed multi epitope vaccine. Here, T = Residue of TLR-4, while V = Residue of Vaccine Candidate. hb = Hydrogen Bond

| Amino Acid Residue Involved | Interacting Amino Acid Residue(s) | Distance | Interaction Type |
| --- | --- | --- | --- |
| T:35:Asn | V:131:Arg | 2.9 Å | 1x hb |
| T:39:Gln | V:92:Arg | 2.7 Å | 1x hb |
|  | V:100:Ala | 2.7 Å | 1x hb |
| T:41:Met | V:92:Arg | 2.7 Å | 1x hb |
| T:51:Asn | V:149:Tyr | 3.0 Å | 1x hb |
| T:264:Arg | V:45:Ala | 3.0 Å | 1x hb |
| T:433:Asn | V:58:Asp | 2.6 Å | 1x hb |
| T:458:His | V:58:Asp | 2.6 Å | 1x hb |
|  | V:60:Asp | 3.1 Å | 1x hb |
| T:507:Gln | V:61:Phe | 2.7 Å | 1x hb |
| T:529:His | V:67:Phe | 3.4 Å | 1x hb, 1x pi stack |
| T:573:Phe | V:200:His | 3.2 Å | 1x pi stack |
| T:578:Gln | V:68:Gly | 3.3 Å | 1x hb |
| T:598:Arg | V:197:His | 2.8 Å | 1x hb |
| T:603:Glu | V:75:Tyr | 2.6 Å | 1x hb |
|  | V:79:Lys | 2.6 Å | 1x hb, 1x salt bridge |
| V:45:Ala | T:264:Arg | 3.0 Å | 1x hb |
| V:58:Asp | T:433:Asn | 2.6 Å | 1x hb |
|  | T:458:His | 2.6 Å | 1x hb |
| V:60:Asp | T:458:His | 3.1 Å | 1x hb |
| V:61:Phe | T:507:Gln | 2.7 Å | 1x hb |
| V:64:Ala | T:505:Gln | 2.7 Å | 1x hb |
|  | T:529:His | 3.7 Å | 1x hb |
| V:65:Ala | T:529:His | 2.6 Å | 1x hb |
|  | T:553:Leu | 4.0 Å | 1x hb |
| V:67:Phe | T:529:His | 3.4 Å | 1x pi stack |
| V:68:Gly | T:578:Gln | 3.3 Å | 1x hb |
| V:75:Tyr | T:603:Glu | 2.6 Å | 1x hb |
| V:79:Lys | T:603:Glu | 2.6 Å | 1x hb, 1x salt bridge |
| V:100:Ala | T:39:Gln | 3.4 Å | 1x hb |
| V:131:Arg | T:35:Asn | 2.9 Å | 1x hb |
| V:149:Tyr | T:51:Asn | 3.0 Å | 1x hb |
| V:197:His | T:598:Arg | 2.8 Å | 1x hb |
| V:200:His | T:573:Phe | 3.2 Å | 1x pi stack |
